# Modeling Linkage Disequilibrium Increases Accuracy of Polygenic Risk Scores

**DOI:** 10.1101/015859

**Authors:** Bjarni J. Vilhjálmsson, Jian Yang, Hilary Finucane, Alexander Gusev, Sara Lindström, Stephan Ripke, Giulio Genovese, Po-Ru Loh, Gaurav Bhatia, Ron Do, Tristan Hayeck, Hong-Hee Won, Schizophrenia Working Group of the Psychiatric Genomics Consortium, the Discovery, Biology, Risk of Inherited Variants in Breast Cancer (DRIVE) study, Sekar Kathiresan, Michele Pato, Carlos Pato, Rulla Tamimi, Eli Stahl, Noah Zaitlen, Bogdan Pasaniuc, Mikkel H. Schierup, Philip De Jager, Nikolaos A. Patsopoulos, Steve McCarroll, Mark Daly, Shaun Purcell, Daniel Chasman, Benjamin Neale, Michael Goddard, Peter Visscher, Peter Kraft, Nick Patterson, Alkes L. Price

## Abstract

Polygenic risk scores have shown great promise in predicting complex disease risk, and will become more accurate as training sample sizes increase. The standard approach for calculating risk scores involves LD-pruning markers and applying a *P*-value threshold to association statistics, but this discards information and may reduce predictive accuracy. We introduce a new method, LDpred, which infers the posterior mean causal effect size of each marker using a prior on effect sizes and LD information from an external reference panel. Theory and simulations show that LDpred outperforms the pruning/thresholding approach, particularly at large sample sizes. Accordingly, prediction *R*^2^ increased from 20.1% to 25.3% in a large schizophrenia data set and from 9.8% to 12.0% in a large multiple sclerosis data set. A similar relative improvement in accuracy was observed for three additional large disease data sets and when predicting in non-European schizophrenia samples. The advantage of LDpred over existing methods will grow as sample sizes increase.

## Introduction

Polygenic risk scores (PRS) computed from genome-wide association study (GWAS)summary statistics have proven valuable for predicting disease risk and understanding the genetic architecture of complex traits. PRS were used to predict genetic risk in a schizophrenia GWAS for which there was only one genome-wide significant locus^1^ and have been widely used to predict genetic risk for many traits^1-15^. PRS can also be used to draw inferences about genetic architectures within and across traits^12,13,16-18^. As GWAS sample sizes grow the prediction accuracy of PRS will increase and may eventually yield clinically actionable predictions^16,19-21^. However, as noted in recent work^19^, current PRS methods do not account for effects of linkage disequilibrium (LD), which limits their predictive value, especially for large samples. Indeed, our simulations show that, in the presence of LD, the prediction accuracy of the widely used approach of LD-pruning followed by *P*-value thresholding^1,6,8,9,12,13,15,16,19,20^ falls short of the heritability explained by the SNPs (**Figure 1** and **Supplementary Figure 1**; see Online Methods).

**Figure 1:**
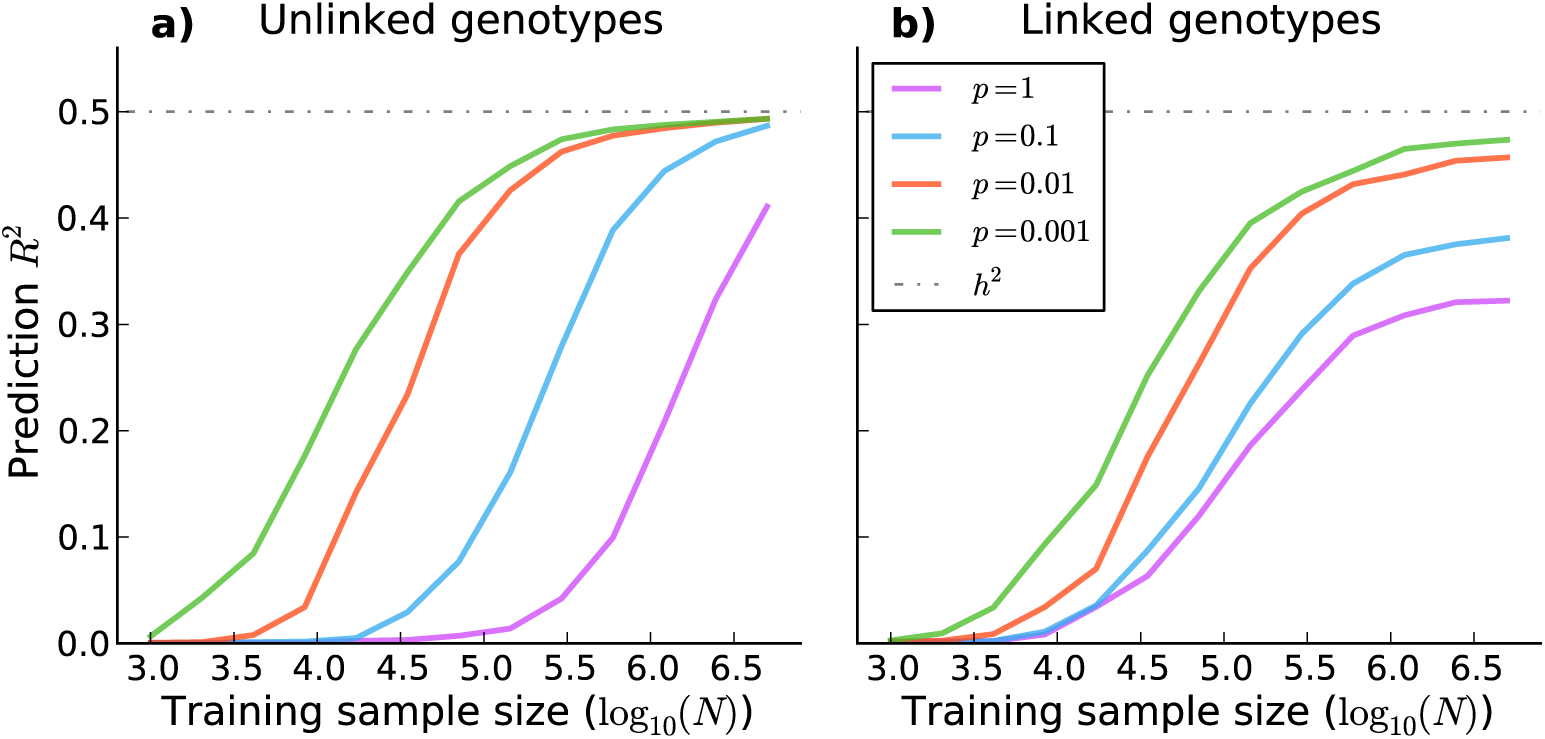
The performance of polygenic risk scores using LD-pruning (*r*^2^<0.2) followed by thresholding (P+T) with optimized threshold when applied to simulated genotypes with and without LD. The prediction accuracy, as measured by squared correlation between the true phenotypes and the polygenic risk scores (prediction *R*^2^), is plotted as a function of the training sample size. The results are averaged over 2000 simulated traits with 200K simulated genotypes where the fraction of causal variants *p* was let vary. In **a)** the simulated genotypes are unlinked. In **b)** the simulated genotypes are linked, where we simulated independent batches of 100 markers where the squared correlation between adjacent variants in a batch was fixed to 0.9.

One possible solution to this problem is to use one of the many available prediction methods that require genotype data as input, including genomic BLUP—which assumes an infinitesimal distribution of effect sizes—and its extensions to non-infinitesimal mixture priors^22-28^. However, these methods are not applicable to GWAS summary statistics when genotype data are unavailable due to privacy concerns or logistical constraints, as is often the case. In addition, many of these methods become computationally intractable at the very large sample sizes (>100K individuals) that would be required to achieve clinically relevant predictions for most common diseases^16,19,20^.

In this study we propose a Bayesian polygenic risk score, LDpred, which estimates posterior mean causal effect sizes from GWAS summary statistics assuming a prior for the genetic architecture and LD information from a reference panel. By using a point-normal mixture prior^26,29^ for the marker effects, LDpred can be applied to traits and diseases with a wide range of genetic architectures. Unlike LD-pruning followed by *P*-value thresholding, LDpred has the desirable property that its prediction accuracy converges to the heritability explained by the SNPs as sample size grows (see below). Using simulations based on real genotypes we compare the prediction accuracy of LDpred to the widely used approach of LD-pruning followed by *P*-value thresholding^1,6,8,9,12,13,15,16,19,20,30^, as well as other approaches that train on GWAS summary statistics. We apply LDpred to seven common diseases for which raw genotypes are available in small sample size, and to five common diseases for which only summary statistics are available in large sample size.

## Results

### Overview of Methods

LDpred calculates the posterior mean effects from GWAS summary statistics conditional on a genetic architecture prior and LD information from a reference panel. The inner product of these re-weighted effect sizes with test sample genotypes is the posterior mean phenotype and thus, under the model assumptions and available data, the best unbiased prediction (see Online Methods). The prior for the effect sizes is a point-normal mixture distribution, which allows for non-infinitesimal genetic architectures. The prior has two parameters, the heritability explained by the genotypes, and the fraction of causal markers (i.e. the fraction of markers with non-zero effects). The heritability parameter is estimated from GWAS summary statistics, accounting for sampling noise and LD^31-33^ (see Online Methods). The fraction of causal markers is allowed to vary and can be optimized with respect to prediction accuracy in a validation data set, analogous to how *P*-value thresholds are varied in standard PRS. We approximate LD using data from a reference panel (e.g. independent validation data). The posterior mean effect sizes are estimated via Markov Chain Monte Carlo (MCMC), and applied to validation data to obtain polygenic risk scores. In the special case of no LD, posterior mean effect sizes with a point-normal prior can be viewed as a soft threshold, and can be computed analytically (**Supplementary Figure 2**; see Online Methods). We have released open-source software implementing the method (see Web Resources).

A key feature of LDpred is that it relies on GWAS summary statistics, which are often available even when raw genotypes are not. In our comparison of methods we therefore focus primarily on polygenic risk scores that rely on GWAS summary statistics. The main approaches that we compare LDpred with are listed in **Table 1**. These include Polygenic Risk Score using all markers (PRS-all), LD-pruning followed by *P*-value thresholding (P+T) and LDpred specialized to an infinitesimal prior (LDpred-inf) (see Online Methods). We note that LDpred-inf is an analytic method, since posterior mean effects are closely approximated by:

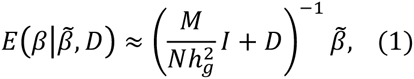

**Table 1:**
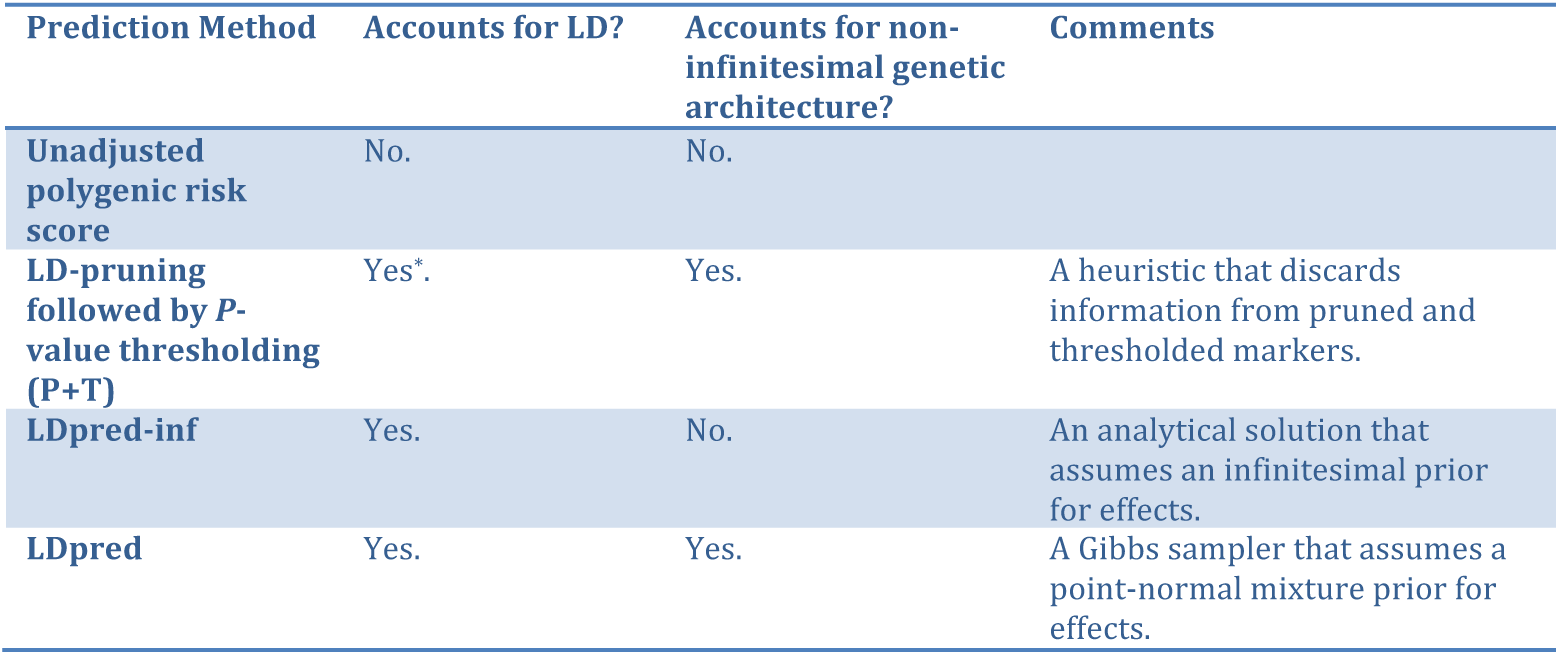
A list of the main polygenic risk score methods (using summary association statistics as input) considered in this study. (*Although P+T prunes SNPs in high LD, it ignores bias induced by linked causal markers.)

| Prediction Method | Accounts for LD? | Accounts for non-infinitesimal genetic architecture? | Comments |
| --- | --- | --- | --- |
| Unadjusted polygenic risk score | No. | No. |  |
| LD-pruning followed by *P*-value thresholding (P+T) | Yes*. | Yes. | A heuristic that discards information from pruned and thresholded markers. |
| LDpred-inf | Yes. | No. | An analytical solution that assumes an infinitesimal prior for effects. |
| LDpred | Yes. | Yes. | A Gibbs sampler that assumes a point-normal mixture prior for effects. |

where *D* denotes the LD matrix between the markers in the training data and 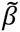 denotes the marginally estimated marker effects (see Online Methods). LDpred-inf (using GWAS summary statistics) is analogous to genomic BLUP (using raw genotypes), as it assumes the same prior.

### Simulations

We first considered simulations with simulated genotypes (see Online Methods). Accuracy was assessed using squared correlation (prediction *R*^2^) between observed and predicted phenotype. The Bayesian shrink imposed by LDpred generally performed well in simulations without LD (**Supplementary Figure 3**); in this case, posterior mean effect sizes can be obtained analytically (see Online Methods). However, LDpred performed particularly well in simulations with LD (**Supplementary Figure 4**); the larger improvement (e.g. vs. P+T) in this case indicates that the main advantage of LDpred is in its explicit modeling of LD. Simulations under a Laplace mixture distribution prior gave similar results (see **Supplementary Figure 5**). Below we focus on simulations with real Wellcome Trust Case Control Consortium genotypes, which have more realistic LD properties.

Using real Wellcome Trust Case Control Consortium (WTCCC) genotypes^34^ (15,835 samples and 376,901 markers, after QC), we simulated infinitesimal traits with heritability set to 0.5 (see Online Methods). We extrapolated results for larger sample sizes (*N_eff_*) by restricting the simulations to a subset of the genome (smaller *M*), leading to larger *N/M*. Results are displayed in **Figure 2a**. LDpred-inf and LDpred (which are expected to be equivalent in the infinitesimal case) performed well in these simulations—particularly at large values of *N_eff_*, consistent with the intuition from Equation (1) that the LD adjustment arising from the reference panel LD matrix (*D*) is more important when 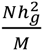 is large. On the other hand, P+T performs less well, consistent with the intuition that pruning markers loses information.

**Figure 2:**
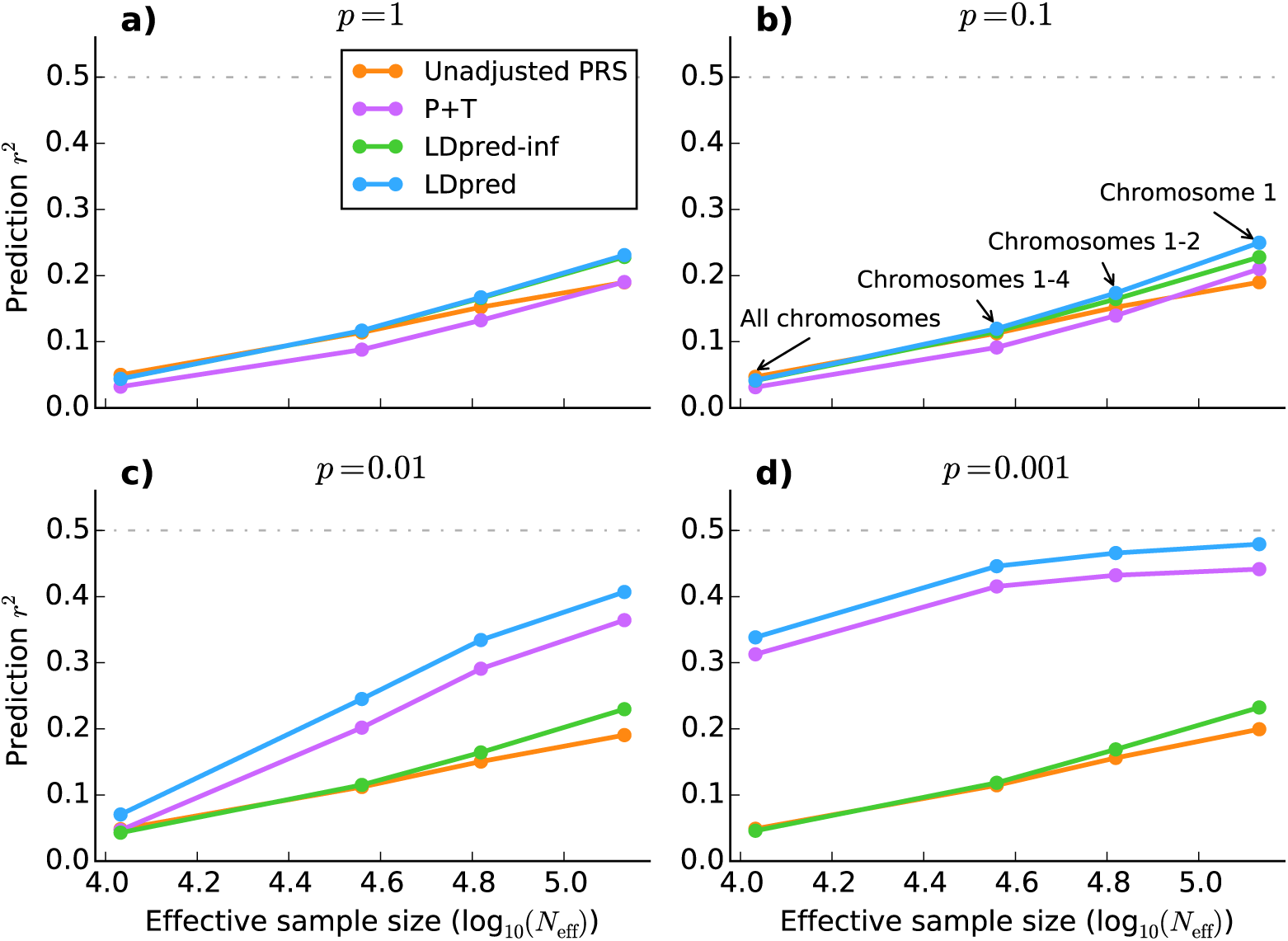
Comparison between the four different methods listed in Table 1 when applied to simulated traits with WTCCC genotypes. The four subfigures **a-d**, correspond to different values of the fraction of simulated causal markers (*p*) with (non-zero) effect sizes sampled from a Gaussian distribution. To aid interpretation of the results, we plot the accuracy against the effective sample size defined as 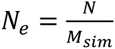, where *N*=10,786 is the training sample size, *M*=376,901 is the total number of SNPs, and *M_sim_* is the actual number of SNPs used in each simulation: 376,901 (all chromosomes), 112,185 (chromosomes 1-4), 61,689 (chromosomes 1-2) and 30,004 (chromosome 1), respectively. The effective sample size is the sample size that maintains the same N/M ratio if using all SNPs.

We next simulated non-infinitesimal traits using real WTCCC genotypes, varying the proportion *p* of causal markers (see Online Methods). Results are displayed in **Figure 2b-d**. LDpred outperformed all other approaches including P+T, particularly at large values of *N/M*. For *p*=0.01 and *p*=0.001, the methods that do not account for non-infinitesimal architectures (Unadjusted PRS and LDpred-inf) perform poorly, and P+T is second best among these methods. Comparisons to additional methods are provided in **Supplementary Figure 6**; in particular, LDpred outperforms other recently proposed approaches that use LD from a reference panel^14,35^.

Besides accuracy (prediction *R*^2^), another measure of interest is calibration. A predictor is correctly calibrated if a regression of the true phenotype vs. the predictor yields a slope of 1, and mis-calibrated otherwise; calibration is particularly important for risk prediction in clinical settings. In general, unadjusted PRS and P+T yield poorly calibrated risk scores. On the other hand, the Bayesian approach provides correctly calibrated predictions (if the prior accurately models the true genetic architecture and the LD is appropriately accounted for), avoiding the need for re-calibration at the validation stage. The calibration slopes for the simulations using WTCCC genotypes are given in **Supplementary Figure 7**. As expected, LDpred provides much better calibration than other approaches.

### Application to WTCCC disease data sets

We compared LDpred to other summary statistic based methods across the 7 WTCCC disease data sets^34^, using 5-fold cross validation (see Online Methods). Results are displayed in **Figure 3**. (We used Nagelkerke *R*^2^ as our primary figure of merit in order to be consistent with previous work^1,9,13,15^, but we also provide results for observed-scale *R*^2^, liability-scale *R*^2^ [ref. ^36^] and AUC^37^ in **Supplementary Table 1**; the relationship between these metrics is discussed in Online Methods).

**Figure 3:**
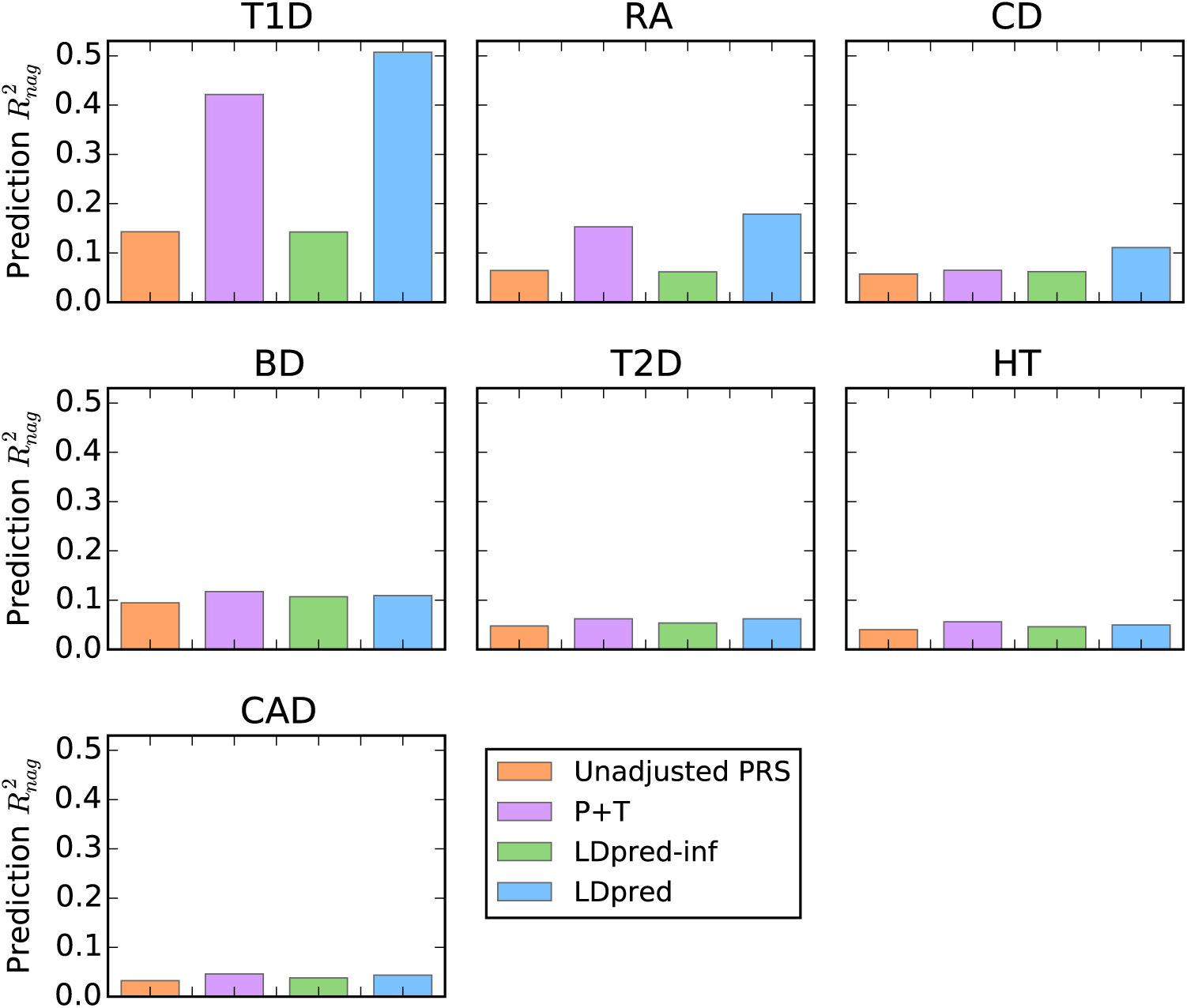
Comparison of methods when applied to 7 WTCCC disease data sets, type-1 diabetes (T1D), rheumatoid arthritis (RA), Chron’s disease (CD), bipolar disease (1D), type-2 diabetes (T2D), hypertension (HT), coronary artery disease (CAD). The Nagelkerke prediction *R*^2^ is shown on the y-axis, see **Supplementary Table 1** for other metrics. LDpred significantly improved the prediction accuracy for the immune-related diseases T1D, RA, and CD (see main text).

LDpred attained significant improvement in prediction accuracy over P+T for T1D (*P*-value=4.4e-15), RA (*P*-value=1.2e-5), and CD (*P*-value=2.7e-8), similar to previous results on the same data using BSLMM^27^. For these three immune-related disorders the MHC region explains a large amount of the overall variance, representing an extreme special case of a non-infinitesimal genetic architecture. We note that LDpred and BSLMM both explicitly model non-infinitesimal architectures; however, unlike LDpred, BSLMM requires full genotype data and cannot be applied to large summary statistic data sets (see below). For the other diseases with more complex genetic architectures the prediction accuracy of LDpred was similar to P+T, potentially due to insufficient training sample size for modeling LD to have a large impact. The inferred heritability parameters and optimal *p* parameters for LDpred, as well as the optimal thresholding parameters for P+T, are provided in **Supplementary Table 2**. The calibration of the predictions for the different approaches is shown in **Supplementary Table 3** Consistent with our simulations, LDpred provides much better calibration than other approaches.

### Application to five large summary statistic data sets

We applied LDpred to five diseases—schizophrenia (SCZ), multiple sclerosis (MS), breast cancer (BC), type 2 diabetes (T2D) and coronary artery disease (CAD)—for which we had GWAS summary statistics for large sample sizes (ranging from 27K to 86K individuals) and raw genotypes for an independent validation data set (see Online Methods). Prediction accuracies for LDpred and other methods are reported in **Figure 4** (Nagelkerke *R*^2^) and **Supplementary Table 4** (other metrics).

**Figure 4:**
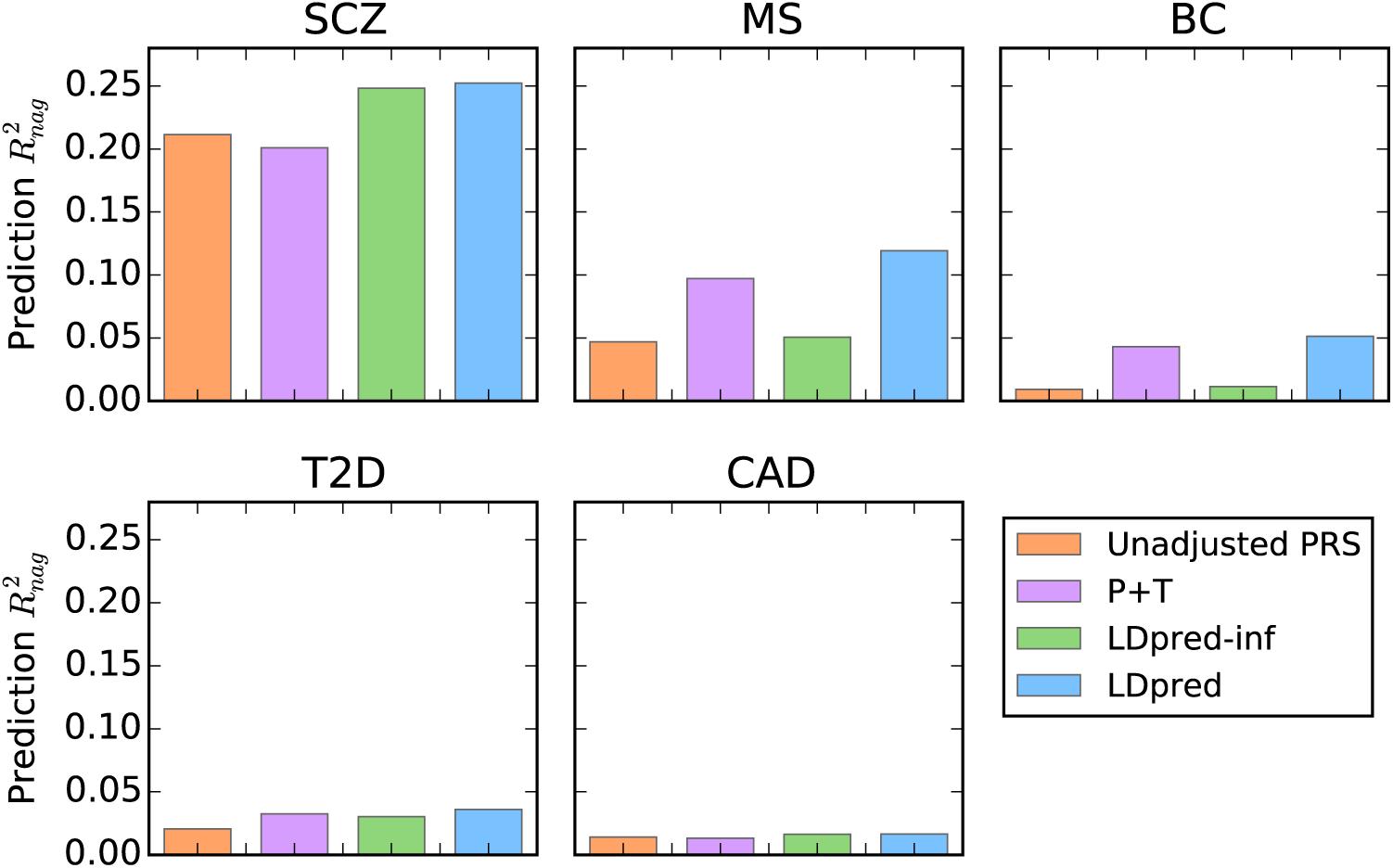
Comparison of prediction accuracy for 5 different diseases, schizophrenia (SCZ), multiple sclerosis (MS), breast cancer (RC), type-2 diabetes (T2D), and coronary artery disease (CAD). The risk scores were trained using large GWAS summary statistics data sets and used to predict in independent validation data sets. The Nagelkerke prediction *R*^2^ is shown on the y-axis (see **Supplementary Table 1** for other metrics). LDpred improved the prediction *R*^2^ by 11-25% compared to LD-pruning + Thresholding (P+T). SCZ results are shown for the SCZ-MGS validation cohort used in recent studies^9,13,15^, but LDpred also produced a large improvement for the independent SCZ-ISC validation cohort (**Supplementary Table 4)**.

For all 5 diseases, LDpred provided significantly better predictions than other approaches (for the improvement over P+T the *P-*values were 6.3e-47 for SCZ, 2.0e-14 for MS, 0.020 for BC, 0.004 for T2D, and 0.017 for CAD). The relative increase in Nagelkerke *R*^2^ over other approaches ranged from 11% for T2D to >25% for SCZ. This is consistent with our simulations showing larger improvements when the trait is highly polygenic, as is known to be the case for SCZ^15^. We note that for both CAD and T2D, the accuracy attained using >60K training samples from large meta-analyses (**Figure 4**) is actually lower than the accuracy attained using <5K training samples from WTCCC (**Figure 3**). This result is independent of the prediction method applied, and demonstrates the challenges of potential heterogeneity in large meta-analyses (although prediction results based on cross-validation in a single cohort should be viewed with caution^20^).

Parameters inferred by LDpred and other methods are provided in **Supplementary Table 5**, and calibration results are provided in **Supplementary Table 6**, with LDpred again attaining the best calibration. Finally, we applied LDpred to predict SCZ risk in non-European validation samples of both African and Asian descent (see Online Methods). Although prediction accuracies were lower in absolute terms, we observed similar relative improvements for LDpred vs. other methods (**Supplementary Tables 7** **and** **8**).

## Discussion

Polygenic risk scores are likely to become clinically useful as GWAS sample sizes continue to grow^16,19^. However, unless LD is appropriately modeled, their predictive accuracy will fall short of their maximal potential. Our results show that LDpred is able to address this problem—even when only summary statistics are available—by estimating posterior mean effect sizes using a point-normal prior and LD information from a reference panel. Intuitively, there are two reasons for the relative gain in prediction accuracy of LDpred polygenic risk scores over LD-pruning followed by *P*-value thresholding (P+T). First, LD-pruning discards informative markers, and thereby limits the overall heritability explained by the markers. Second, LDpred accounts for the effects of linked markers, which can otherwise lead to biased estimates. These limitations hinder P+T regardless of the LD-pruning and *P*-value thresholds used.

Although we focus here on methods that only require summary statistics, we note the parallel advances that have been made in methods that require raw genotypes^23,25-28,38-40^ as training data. Some of those methods employ a Variational Bayes (Iterative Conditional Expectation) approach to reduce their running time^25,26,38,40^ (and report that results are similar to MCMC^40^), but we found that MCMC generally obtains more robust results than Variational Bayes when analyzing summary statistics, perhaps because the LD information is only approximate. Our use of a point-normal mixture prior is consistent with some of those studies^26^, although different priors were used by other studies, e.g. a mixture of normals^24,27^. One recent study proposed an elegant approach for handling case-control ascertainment while including genome-wide significant associations as fixed effects^39^; however, the correlations between distal causal SNPs induced by case-control ascertainment do not impact summary statistics from marginal analyses, and explicit modeling of non-infinitesimal effect size distributions will appropriately avoid shrinking genome-wide significant associations (**Supplementary Figure 2**).

While LDpred is a substantial improvement on existing methods for conducting polygenic prediction using summary statistics, it still has limitations. First, the method’s reliance on LD information from a reference panel requires that the reference panel be a good match for the population from which summary statistics were obtained; in the case of a mismatch, prediction accuracy may be compromised. One potential solution is the broad sharing of summary LD statistics, which has previously been advocated in other settings^41^. Second, the point-normal mixture prior distribution used by LDpred may not accurately model the true genetic architecture, and it is possible that other prior distributions may perform better in some settings. Third, in those instances where raw genotypes are available, fitting all markers simultaneously (if computationally tractable) may achieve higher accuracy than methods based on marginal summary statistics. Fourth, as with other prediction methods, heterogeneity across cohorts may hinder prediction accuracy; our results suggest that this could be a major concern in some data sets. Fifth, joint analysis of multiple traits—which can potentially increase prediction accuracy—is not considered here, and remains as a future direction^42^. Sixth, we assume that summary statistics have been appropriately corrected for genetic ancestry, but if this is not the case then the prediction accuracy may be misinterpreted^20^, or may even decrease^43^. Seventh, our analyses have focused on common variants; LD reference panels are likely to be inadequate for rare variants, motivating future work on how to treat rare variants in polygenic risk scores. Finally, we have not considered the advantages of different prior distributions across genomic regions^28^ or functional annotation classes^44^, whose incorporation into methods for polygenic prediction remains as a future direction. Despite these limitations, LDpred is likely to be broadly useful in leveraging summary statistic data sets for polygenic prediction.

## Online Methods

### Phenotype model

Let *Y* be a N×1 phenotype vector and *X* a *N*×*M* genotype matrix, where the *N* is the number of individuals and *M* is the number of genetic variants. For simplicity, we will assume throughout that the phenotype *Y* and individual genetic variants *X_i_* have been mean-centered and standardized to have variance 1. We model the phenotype as a linear combination of *M* genetic effects and an independent environmental effect *ε*, i.e. 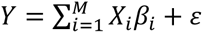, where *X_i_* denotes the *i*’th genetic variant, *β_i_* its true effect, and E the environmental and noise contribution. In this setting the (marginal) least square estimate of an individual marker effect is 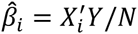. For clarity we implicitly assume that we have the standardized effect estimates available to us as summary statistics. In practice, we usually have other summary statistics, including the *P*-value and direction of the effect estimates, from which we infer the standardized effect estimates. First, we exclude all markers with ambiguous effect directions, i.e. A/T and G/C SNPs. Second, from the *P*-values we obtain Z-scores, and multiply them by the sign of the effects (obtained from the effect estimates or effect direction). Finally we approximate the least square estimate for the effect by 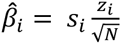, where *s_i_* is the sign, and *z_i_* is the Z-score as obtained from the *P*-value. If the trait is a case control trait, this transformation from the *P*-value to the effect size can be thought of as being an effect estimate for an underlying quantitative liability or risk trait^45^.

### Polygenic risk score using all markers (PRS-all)

The polygenic risk score using all genotyped markers is simply the sum of all the estimated marker effects for each allele, i.e. the standard unadjusted polygenic score for the *i*^th^ individual is 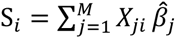.

### LD-pruning followed by thresholding (P+T)

In practice, the prediction accuracy is improved if the markers are LD-pruned and *P-*value pruned a priori. Informed LD-pruning (also known as LD-clumping), which preferentially prunes the less significant marker, often yields much more accurate predictions than pruning random markers. Applying a *P*-value threshold, i.e. only markers that achieve a given significance thresholds are used, also improves prediction accuracies for many traits and diseases. In this paper the LD-pruning followed by thresholding approach refers to the strategy of first applying informed LD-pruning with *r^2^* threshold of 0.2, and subsequently *P*-value thresholding where the *P*-value threshold is optimized over a grid with respect to prediction accuracy in the validation data.

### Bayesian approach in the special case of no LD (Bpred)

Under a model, the optimal linear prediction given some statistic is the posterior mean prediction. This prediction is optimal in the sense that it minimizes the prediction error variance and is unbiased in the Bayesian sense^46^. Under the linear model described above, the posterior mean phenotype given GWAS summary statistics and LD is

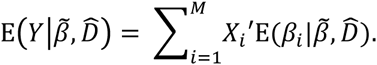

Here 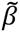 denotes a vector of marginally estimated least square estimates as obtained from the GWAS summary statistics, and 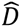 refers to the observed genome-wide LD matrix in the training data, i.e. the samples for which the effect estimates are calculated. Hence the quantity of interest is the posterior mean marker effect given LD information from the GWAS sample and the GWAS summary statistics. In practice we may not have this information available to us and are forced to estimate the LD from a reference panel. In our analysis we used the independent validation data set to estimate the local LD structure in the training data.

The variance of the trait can be partitioned into a heritable part and the noise, i.e. 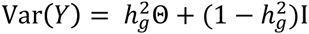, where 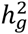 denotes the heritability explained by the genotyped variants, and 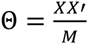 is the SNP-based genetic relationship matrix. We can obtain a trait with the desired covariance structure if we sample the betas independently with mean 0 and variance 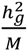. Note that if the effects are independently sampled then this also holds true for correlated genotypes, i.e. when there is LD. However, LD will increase the variance of heritability explained by the genotypes as estimated from the data (due to fewer effective markers).

If we assume that all samples are independent, and that all markers are unlinked and have effects drawn from a Gaussian distribution, i.e. 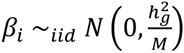. This is an infinitesimal model^47^ where all markers are causal and under it the posterior mean can be derived analytically, as shown by Dudbridge^16^:

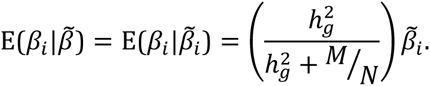

Interestingly, with unlinked markers this infinitesimal shrink factor times the heritability, i.e. 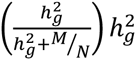, is the expected squared correlation between the polygenic risk score using all (unlinked) markers and the phenotype, regardless of the underlying genetic architecture^48,49^.

An arguably more reasonable prior for the effect sizes is a non-infinitesimal model, where only a fraction of the markers are causal. For this consider the following Gaussian mixture prior

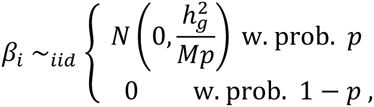

where *p* is the fraction of markers that is causal, is an unknown parameter. Under this model the posterior mean can be derived as (see **Supplementary Note**):

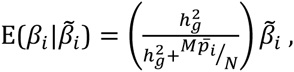

where

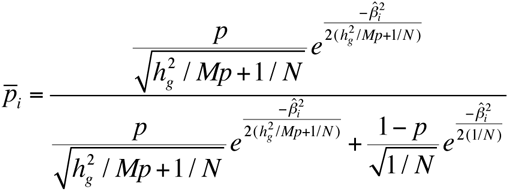

is the posterior probability of an individual marker being causal. In our simulations we refer to this Bayesian shrink without LD as Bpred.

### Bayesian approach in the presence of LD (LDpred)

If we allow for loci to be linked, then we can derive posterior mean effects analytically under a Gaussian infinitesimal prior (described above). We call the resulting method LDpred-inf and it represents a computationally efficient special case of LDpred. If we assume that distant markers are unlinked, the posterior mean for the effect sizes within a small region *l* under an infinitesimal model, is well approximated by

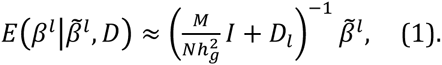

Here *D_l_* denotes the regional LD matrix within the region of LD and 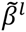 denotes the least square estimated effects within that region. The approximation assumes that the heritability explained by the region is small and that LD with SNPs outside of the region is negligible. Interestingly, under these assumptions the resulting effects approximate the standard mixed model genomic BLUP effects. LDpred-inf is therefore a natural extension of the genomic BLUP to summary statistics. The detailed derivation is given in the **Supplementary Note**. In practice we do not know the LD pattern in the training data, and we need to estimate it using LD in a reference panel.

Deriving an analytical expression for the posterior mean under a non-infinitesimal Gaussian mixture prior is difficult, and we therefore approximate it numerically in LDpred. The Bayesian shrink under the infinitesimal model implies that we can solve it either using a Gauss-Seidel method^50,51^, or via MCMC Gibbs sampling. The Gauss-Seidel method iterates over the markers, and obtains a residual effect estimate after subtracting the effect of neighboring markers in LD. It then applies a univariate Bayesian shrink, i.e. the Bayesian shrink for unlinked markers (described above). It then iterates over the genome multiple times until convergence is achieved. However, we found the Gauss-Seidel approach to be sensitive to model assumptions, i.e. if the LD matrix used differed from the true LD matrix in the training data we observed convergence issues. We therefore decided to use an approximate MCMC Gibbs sampler instead to infer the posterior mean. The approximate Gibbs sampler used by LDpred is similar the Gauss-Seidel approach, except that instead of using the posterior mean to update the effect size, we sample the update from the posterior distribution. Compared to the Gauss-Seidel method this seems to lead to less serious convergence issues. The approximate Gibbs sampler is described in detail in the **Supplementary Note**. To ensure convergence, we shrink the posterior probability of being causal by a fixed factor at each big iteration step *i*, where the shrinkage factor is defined as 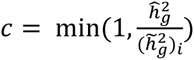, where 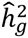 is the estimated heritability using an aggregate approach (see below), and 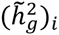 is the estimated genome-wide heritability at each big iteration. To speed upconvergence in the Gibbs-sampler we used Rao-Blackwellization and observed that good convergence was usually attained with less than 100 iterations in practice (see **Supplementary Note**).

#### Estimation of heritability parameter

In the absence of population structure and assuming i.i.d. mean-zero SNP effects, the following equation has been shown to hold

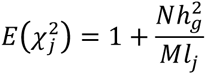

where 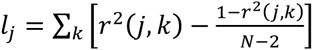, is the LD score for the *j*’th SNP summing over *k* neighboring SNPs in LD. Taking the average of both sides over SNPs and rearranging, we obtain a heritability estimate

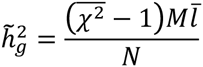

where 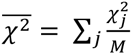, and 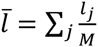. We call this the aggregate estimator, and it is equivalent to LD score regression^31-33^ with intercept constrained to 1 and SNP *j* weighted by 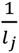. Prediction accuracy is not predicated on the robustness of this estimator, which will be evaluated elsewhere. Following the conversion proposed by Lee *et aL^52^*, we also reported the heritability on the liability scale.

### Simulations

We performed three types of simulations: (1) simulated traits and simulated genotypes; (2) simulated traits, simulated summary statistics and simulated validation genotypes; (3) simulated traits using real genotypes. For most of the simulations we used the point-normal model for effect sizes as described above:

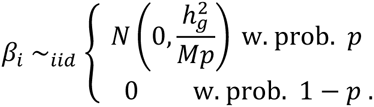

For some of our simulations (**Supplementary Figure 5**) we sampled the non-zero effects from a Laplace distribution instead of a Gaussian distribution. For all of our simulations we used four different values for *p* (the fraction of causal loci). For some of our simulations (**Supplementary Figure 1**) we sampled the *p* parameter from a Beta(*p,1- p*) distribution. The simulated trait was then obtained by summing up the allelic effects for each individual, and adding a Gaussian distributed noise term to fix the heritability. The simulated genotypes were sampled from a standard Gaussian distribution. To emulate linkage disequilibrium (LD) we simulated one genotype or SNP at a time generating batches of 100 correlated SNPs. Each SNP was defined as the sum of the preceding adjacent SNP and some noise, where they were scaled to correspond to a fixed squared correlation between two adjacent SNPs within a batch. We simulated genotypes with the adjacent squared correlation between SNPs set to 0 (unlinked SNPs), and 0.9 when simulating LD.

In order to compare the performance of our method at large sample sizes we simulated summary statistics that we used as training data for the polygenic risk scores. We also simulated a smaller sample (2000 individuals) representing an independent validation data. When there is no LD, the least square effect estimates (summary statistics) are sampled from a Gaussian distribution 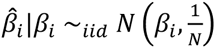, where *β_i_* are the true effects. To simulate marginal effect estimates without genotypes in the presence of LD we first estimate the LD pattern empirically by simulating 100 SNPs for 1000 individuals for a given value (as described above) and average over 1000 simulations. This matrix captures the LD pattern in the validation data since we simulate it using the same procedure (described earlier). Using this LD matrix *D* we then sample the marginal least square estimates within a region of LD (SNP chunk) as 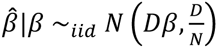, where *D* is the LD matrix.

For the simulations in **Figure 1 b)** and **Supplementary Figures 1**, **3**, and **4**, we simulated least square effect estimates for 200K variants in batches of LD regions with 100 variants each (as described above). We then simulated genotypes for 2000 validation individuals and averaged over 100-500 simulated phenotypes to ensure smooth curves. Depending on the simulation parameters, the actual number of repeats required to achieve a smooth curve varied. For the simulations in **Figure 1 a)** and **Supplementary Figure 2**, we simulated the least square estimates independently by adding an appropriately scaled Gaussian noise term to the true effects.

When simulating traits using the WTCCC genotypes (**Figure 2**) we performed simulations under four different scenarios, representing different number of chromosomes: (1) all chromosomes; (2) chromosomes 1-4; (3) chromosomes 1-2; (4) chromosome 1. We used 16,179 individuals in the WTCCC data, and 376, 901 SNPs that passed quality control. In our simulations we used 3-fold cross validation, using 1/3 of the data as validation data and 2/3 as training data.

### WTCCC Genotype data

We used the Wellcome Trust Case Control Consortium (WTCCC) genotypes^34^ for both simulations and analysis. After quality control, pruning variants with missing rates above 1%, and removing individuals that had genetic relatedness coefficients above 0.05, we were left with 15,835 individuals genotyped for 376,901 SNPs, including 1,819 cases for bipolar disease (BD), 1,862 cases for coronary artery disease (CAD), 1,687 cases for Chron’s disease (CD, 1,907 cases for hypertension (HT), 1,831 cases for rheumatoid arthritis (RA), 1,953 cases for type-1 diabetes (T1D), and 1,909 cases for type-2 diabetes (T2D). For each of the 7 diseases, we performed 5-fold cross-validation on disease cases and 2,867 controls.

### Summary statistics and independent validation data sets

Five large summary statistics data sets were analyzed in this paper. The Psychiatric Genomics Consortium (PGC) 2 schizophrenia summary statistics^15^ consists of 34,241 cases and 45,604 controls. For our purposes we calculated GWAS summary statistics while excluding the ISC (International Schizophrenia Consortium) cohorts and the MGS (Molecular Genetics of Schizophrenia) cohorts respectively. The summary statistics were calculated on a set of 1000 genomes imputed SNPs, resulting in 16.9M statistics. The two independent validation data sets, the ISC and the MGS data sets, both consist of multiple cohorts with individuals of European descent. For both of the validation data sets we used the chip genotypes and filtered individuals with more than 10% of genotype calls missing and filtered SNPs that had more than 1% missing rate and a minor allele frequency greater than 1%. In addition we removed SNPs that had ambiguous nucleotides, i.e. A/T and G/C SNPs. We matched the SNPs between the validation and the GWAS summary statistics data sets based on the SNP *rs*-ID and excluded triplets, SNPs where one nucleotide was unknown, and SNPs that had different nucleotides in different data sets. This was our quality control (QC) procedure for all large summary statistics data sets that we analyzed. After QC, the ISC consisted of 1562 cases and 1994 controls genotyped on 518K SNPs that overlapped with the GWAS summary statistics. The MGS data set consisted of 2681 cases and 2653 controls after QC and had 549K SNPs that overlapped with the GWAS summary statistics.

For multiple sclerosis we used the International Multiple Sclerosis (MS) Genetics Consortium summary statistics^53^. These were calculated with 9,772 cases and 17,376 controls (27,148 individuals in total) for 465K SNPs. As an independent validation data set we used the BWH/MIGEN chip genotypes with 821 cases and 2705 controls^54^. After QC the overlap between the validation genotypes and the summary statistics only consisted of 114K SNPs, which we used for our analysis.

For breast cancer we used the Genetic Associations and Mechanisms in Oncology (GAME-ON) breast cancer GWAS summary statistics, consisting of 16, 003 cases and 41,335 controls (both ER- and ER+ were included in this analysis)^55-58^. These summary statistics were calculated for 2.6M HapMap2 imputed SNPs. As validation genotypes we combined genotypes from five different data sets, BPC3 ER-cases and controls^55^, BRCA NHS2 cases, NHS1 cases and controls from a mammographic density study, CGEMS NHS1 cases^59^, and Kidney Stone NHS2 controls. None of these 307 cases and 560 controls were included in the GWAS summary statistics analysis and thus represent an independent validation data set. We used the chip genotypes that overlapped with the GWAS summary statistics, which resulted in 444K genotypes after QC.

For coronary artery disease we used the transatlantic Coronary ARtery DIsease Genome wide Replication and Meta-analysis (CARDIoGRAM) consortium GWAS summary statistics. These were calculated using 22,233 cases and 64,762 controls (86,995 inviduals in total) for 2.4M SNPs^10^. For the type-2 diabetes we used the DIAbetes Genetics Replication And Meta-analysis (DIAGRAM) consortium GWAS summary statistics. These were calculated using 12,171 cases and 56,862 controls (69,033 individuals in total) for 2,5M SNPs^60^. For both CAD and T2D we used the Womens Genomes Health Study (WGHS) data set as validation data^61^, where we randomly down-sampled the controls. For CAD we validated in 923 cases CVD and 1428 controls, and for T2D we used 1673 cases and 1434 controls. We used the genotyped SNPs that overlapped with the GWAS summary statistics, which amounted to about 290K SNPs for both CAD and T2D after quality control.

For all of these data sets we used the validation data set as an LD-reference for LDpred and when LD-pruning. We calculated risk scores for different *P*-value thresholds using grid values [1E-8, 1E-6, 1E-5, 3E-5, 1E-4, 3E-4, 1E-3, 3E-3, 0.01, 0.03, 0.1, 0.3, 1] and for LDpred we used the mixture probability (fraction of causal markers) values [1E-4, 3E-4, 1E-3, 3E-3, 0.01, 0.03,0.1,0.3,1]. We then reported the optimal prediction value from a validation data for LDpred and P+T respectively.

### Schizophrenia validation data sets with non-European ancestry

For the non-European validation data sets we used the MGS data set as an LD-reference. This required us to coordinate across three different data sets, the GWAS summary statistics, the LD reference genotypes and the validation genotypes. To ensure sufficient overlap of genetic variants across all three data sets we used 1000 genomes imputed MGS genotypes and the 1000 genomes imputed validation genotypes for the three Asian validation data sets (JPN1, TCR1, and HOK2). To limit the number of markers for these data sets we only considered markers that had MAF>0.1. After QC, and removing variants with MAF<0.1, we were left with 1.38 million SNPs and 492 cases and 427 controls in the JPN1 data set, 1.88 million SNPs and 898 cases and 973 controls in the TCR1 data set, and 1.71 million SNPs and 476 cases and 2018 controls in the HOK2 data set.

For the African American validation data set (AFAM) we used the reported GWAS summary statistics data set^15^ to train on. The AFAM data set consisted of 3361 schizophrenia cases and 5076 controls. Since the AFAM data set was not included in that analysis this allowed us to leverage a larger sample size, but at the cost of having fewer SNPs. The overlap between the 1000 genomes imputed MGS genotypes, the HapMap 3 imputed AFAM genotypes and the PGC2 reported summary statistics had 482K SNPs after QC (with a MAF>0.01).

### Prediction accuracy metrics

For simulated quantitative traits, we used squared correlation (*R*^2^). For case-control traits, which include all of the disease data sets analyzed, we used four different metrics. We used Nagelkerke *R*^2^ as our primary figure of merit in order to be consistent with previous work^1,9,13,15^, but also report three other commonly used metrics in **Supplementary Tables 1**, **4**, and **7**: observed scale *R^2^*, liability scale *R^2^*, and the area under the curve (AUC). All of the reported prediction *R^2^* values were adjusted for the top 5 principal components (PCs) in the validation sample (top 3 PCs for breast cancer). The relationship between observed scale *R^2^*, liability scale *R^2^*, and AUC is described in Lee *et al.*^36^. We note that Nagelkerke *R*^2^ is similar to observed scale *R*^2^ (i.e. is also affected by case-control ascertainment), but generally has slightly larger values.

## Web Resources

- LDpred software: http://www.hsph.harvard.edu/alkes-price/software/and http://bitbucket.org/bjarnivilhjalmsson/ldpred
- Genetic Associations and Mechanisms in Oncology (GAME-ON) breast cancer GWAS summary statistics:http://gameon.dfci.harvard.edu
- Type-2 diabetes summary statistics^60^:www.diagram-consortium.org
- Coronary artery disease summary statistics^10^: http://www.cardiogramplusc4d.org
- Schizophrenia summary statistics^15^: http://www.med.unc.edu/pgc/downloads

## Acknowledgments

We thank Shamil Sunayev, Brendan Bulik-Sullivan, Liming Liang, Naomi Wray, Daniel Sorensen, and Esben Agerbo for useful discussions. This research was supported by NIH grants R01 GM105857, R03 CA173785, and U19 CA148065-01. BJV was supported by a Danish Council for Independent Research grant DFF-1325-0014. Members of the Schizophrenia Working Group of the Psychiatric Genomics Consortium and the Discovery, Biology, and Risk of Inherited Variants in Breast Cancer (DRIVE) study are listed in the Supplementary Note. This study made use of data generated by the Wellcome Trust Case Control Consortium (WTCCC) and the Wellcome Trust Sanger Institute. A full list of the investigators who contributed to the generation of the WTCCC data is available at www.wtccc.org.uk. Funding for the WTCCC project was provided by the Wellcome Trust under award 076113

## Supplementary Figures and Tables

**Supplementary Figure 1.**
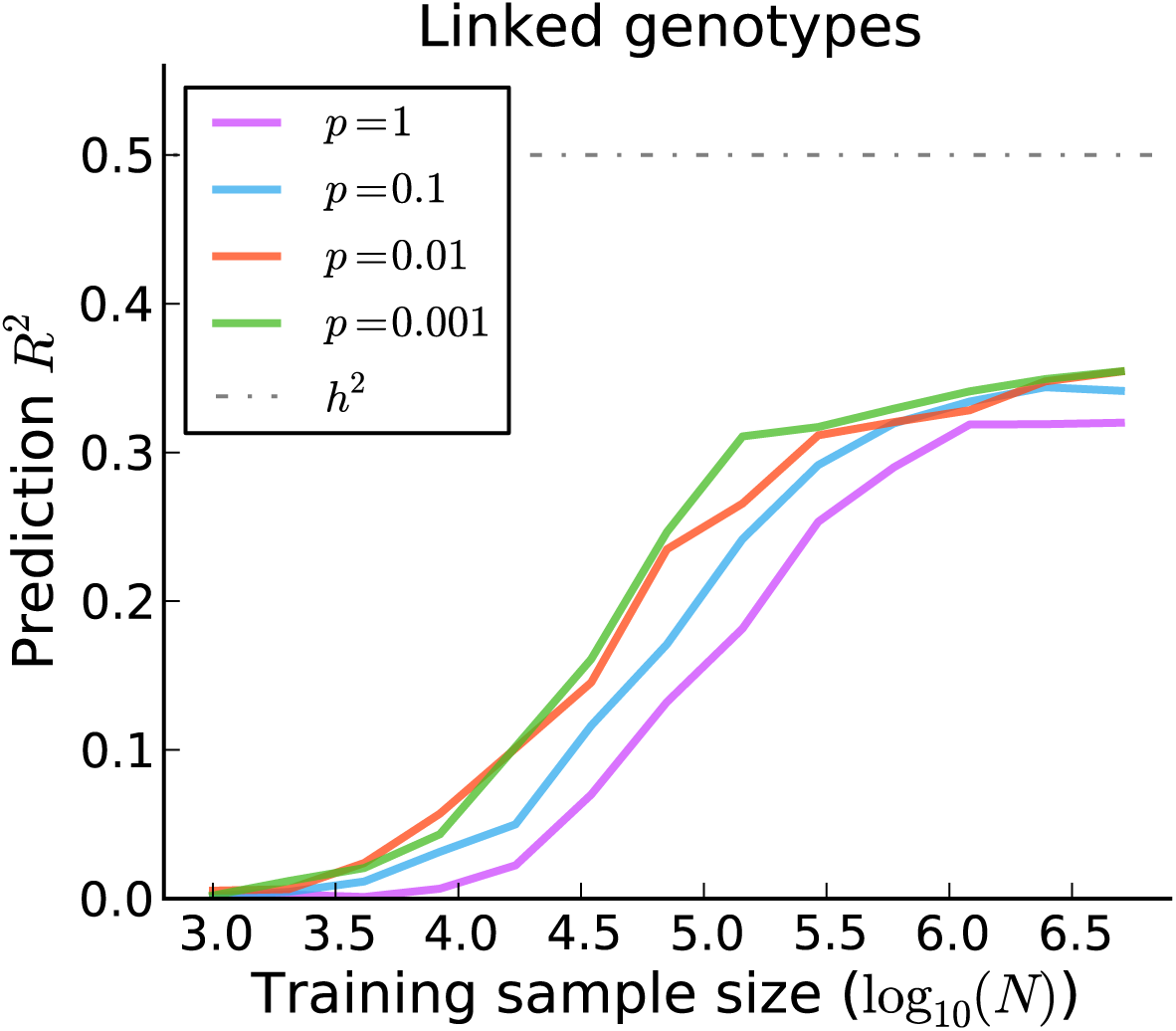
Performance of P+T (LD-pruning followed by thresholding) for an alternative genetic architecture where causal markers cluster. The results are averaged over 3000 simulated traits with 200K simulated genotypes where the average fraction of causal variants *p* was let vary. The simulated genotypes are linked, where we simulated independent batches of 100 markers where the squared correlation between adjacent variants in a batch was fixed to 0.9. For each simulated 100 SNP region of LD, we sampled the *p* parameter from a Beta(*p*, *1-p*) distribution. This will cause causal variants to cluster in some regions of the genome. As expected, the impact of LD on the prediction accuracy of P+T is greater when causal variants cluster, and still substantial for small values of *p*.

**Supplementary Figure 2.**
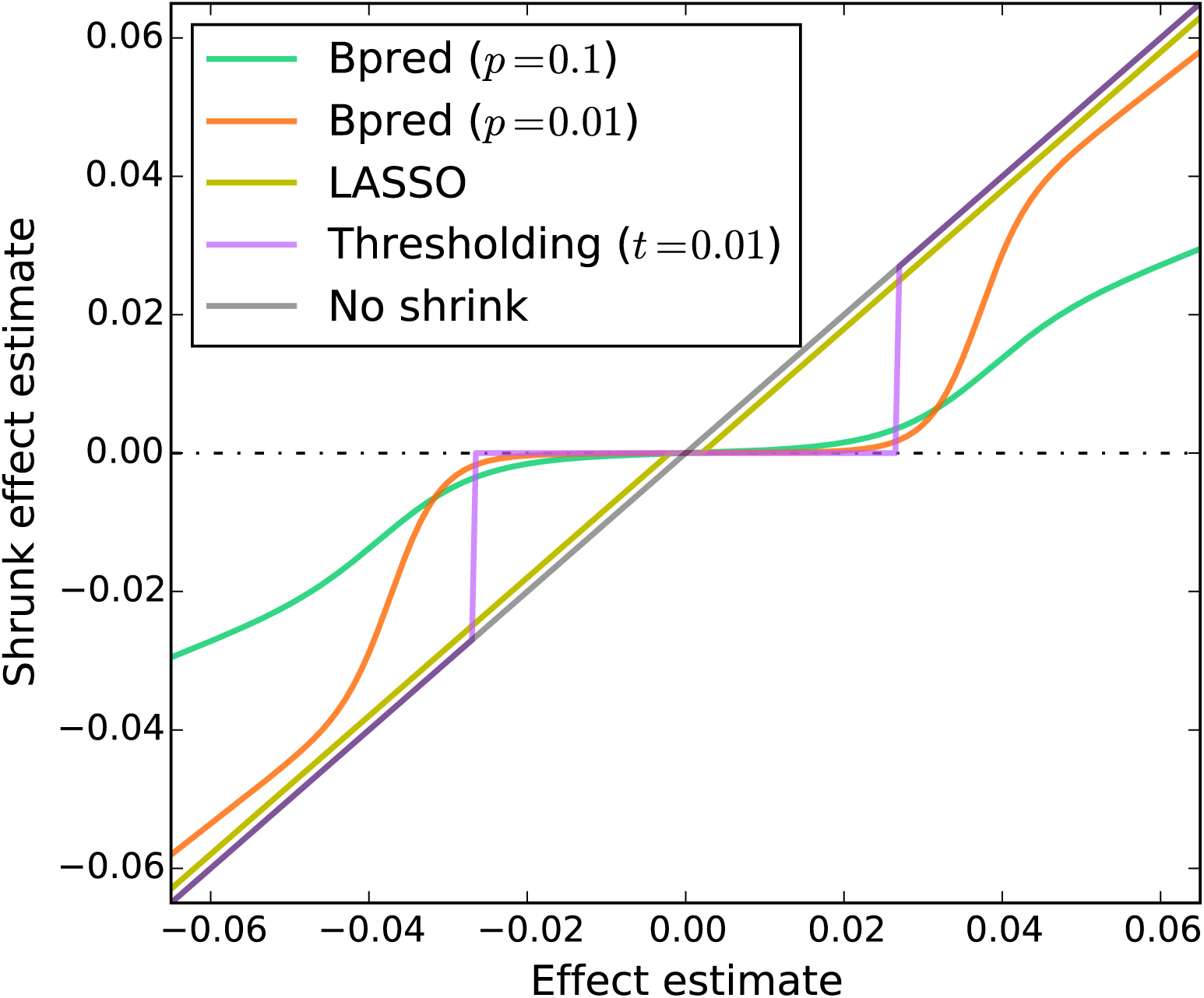
Comparison of five different shrinks in the absence of LD. Bpred corresponds to LDpred without LD and can be derived analytically (see Online Methods and Supplementary Note for details). The marginal (least square) effect estimate is plotted against the shrunk estimate for the five different shrinks. Bpred denotes the analytical solution to LDpred, which can be derived in the absence of LD (see Supplementary Note for details). The Bpred shrink shown here assumes that the heritability is 0.5 and the training sample size is 10,000 and the number of markers is 60,000. Similarly, the LASSO shrink shown here corresponds to the (marginal) posterior mode effect under a Laplace prior for the causal effects. Compared to *P*-value thresholding, and LASSO, Bpred can be viewed as a smoother shrink.

**Supplementary Figure 3.**
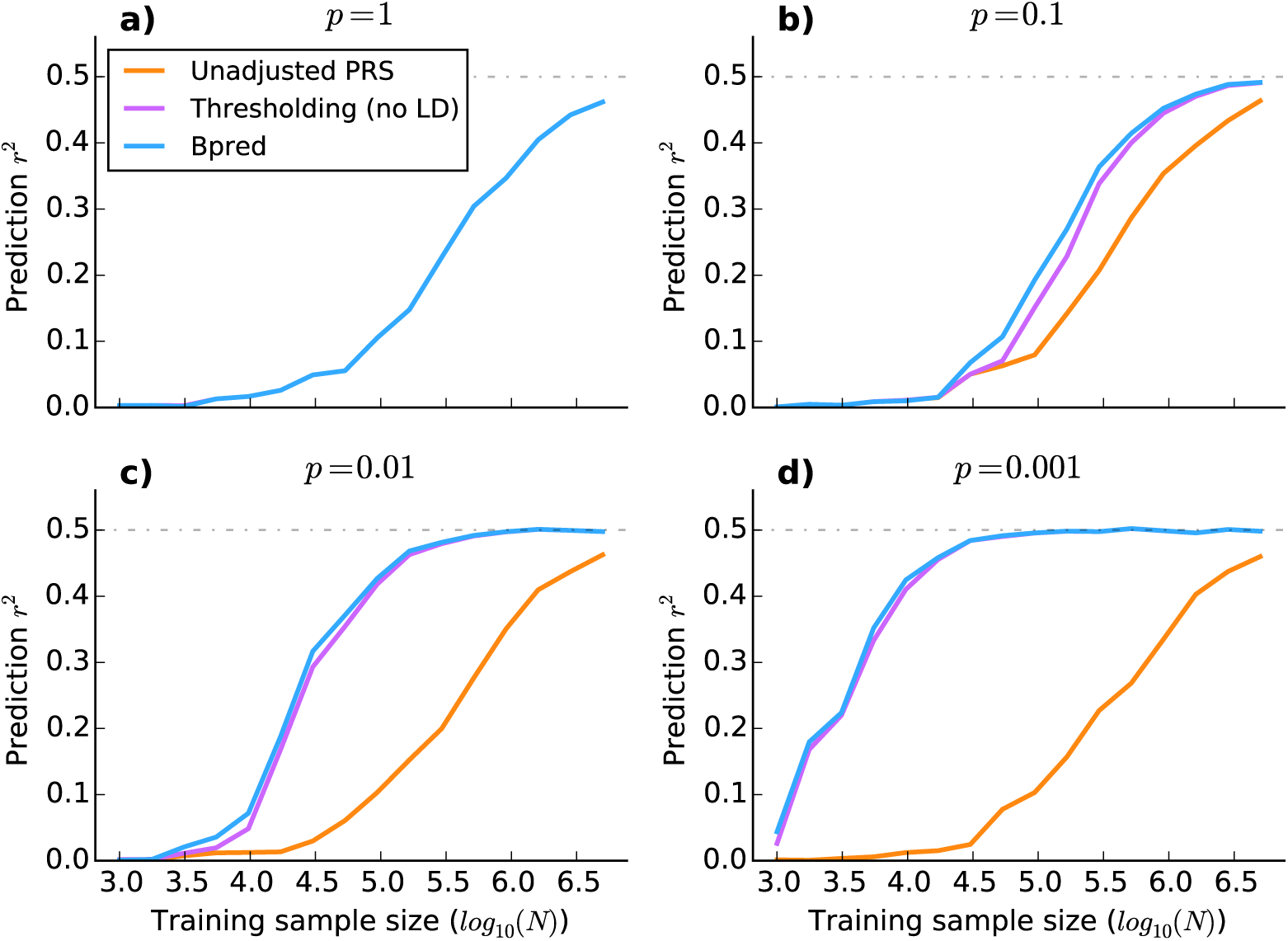
Comparison of methods using simulated genotypes without LD. The four subfigures **a-d** correspond to different genetic architectures where we vary *p*, the fraction of variants with (non-zero) effects drawn from a Gaussian distribution. Bpred denotes the analytical solution to LDpred, which can be derived in the absence of LD (see Supplementary Note for details). As expected, Bpred outperforms *P*-value thresholding in the absence of LD, although not by much.

**Supplementary Figure 4.**
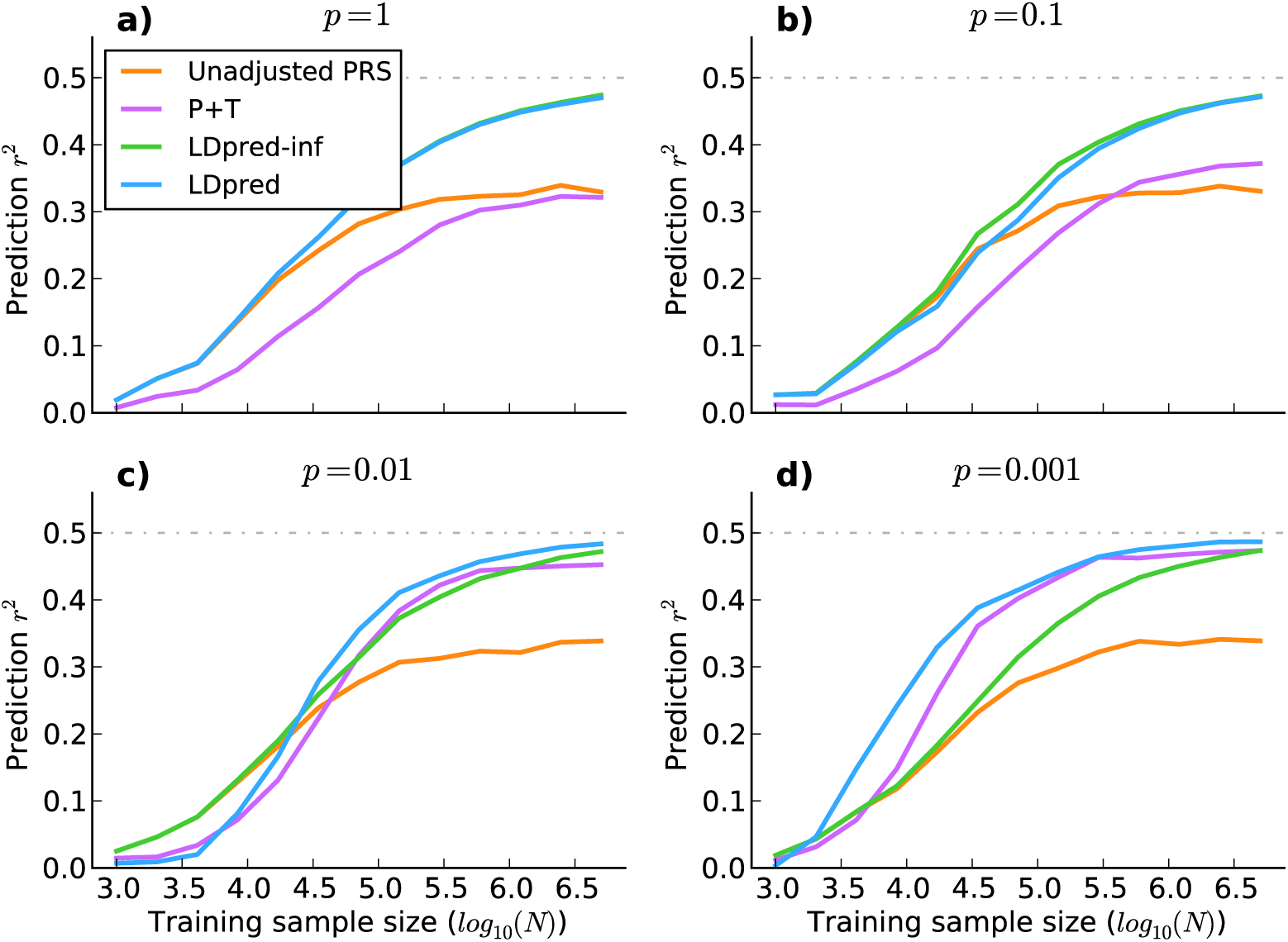
Comparison of methods using simulated genotypes with LD. The four subfigures **a-d** correspond to different genetic architectures where we vary *p*, the fraction of variants with (non-zero) effects drawn from a Gaussian distribution. We simulated marginal least square effect estimates with LD (see Supplementary Note for details). This enabled us to evaluate the behavior of the methods at large sample sizes. The LD structure consisted of 100 SNP regions where adjacent markers had *r^2^=0.9*. For validation we simulated 200000 SNPs in 2000 individuals. For each point in the plot we averaged the results over 2000 independent phenotype simulations keeping the simulated genotypes fixed (see Supplementary Note for details). The behavior of LDpred in the subfigure **c)** for small sample sizes is due to a LD window-size mismatch between the simulated data and the LDpred and P+T methods.

**Supplementary Figure 5.**
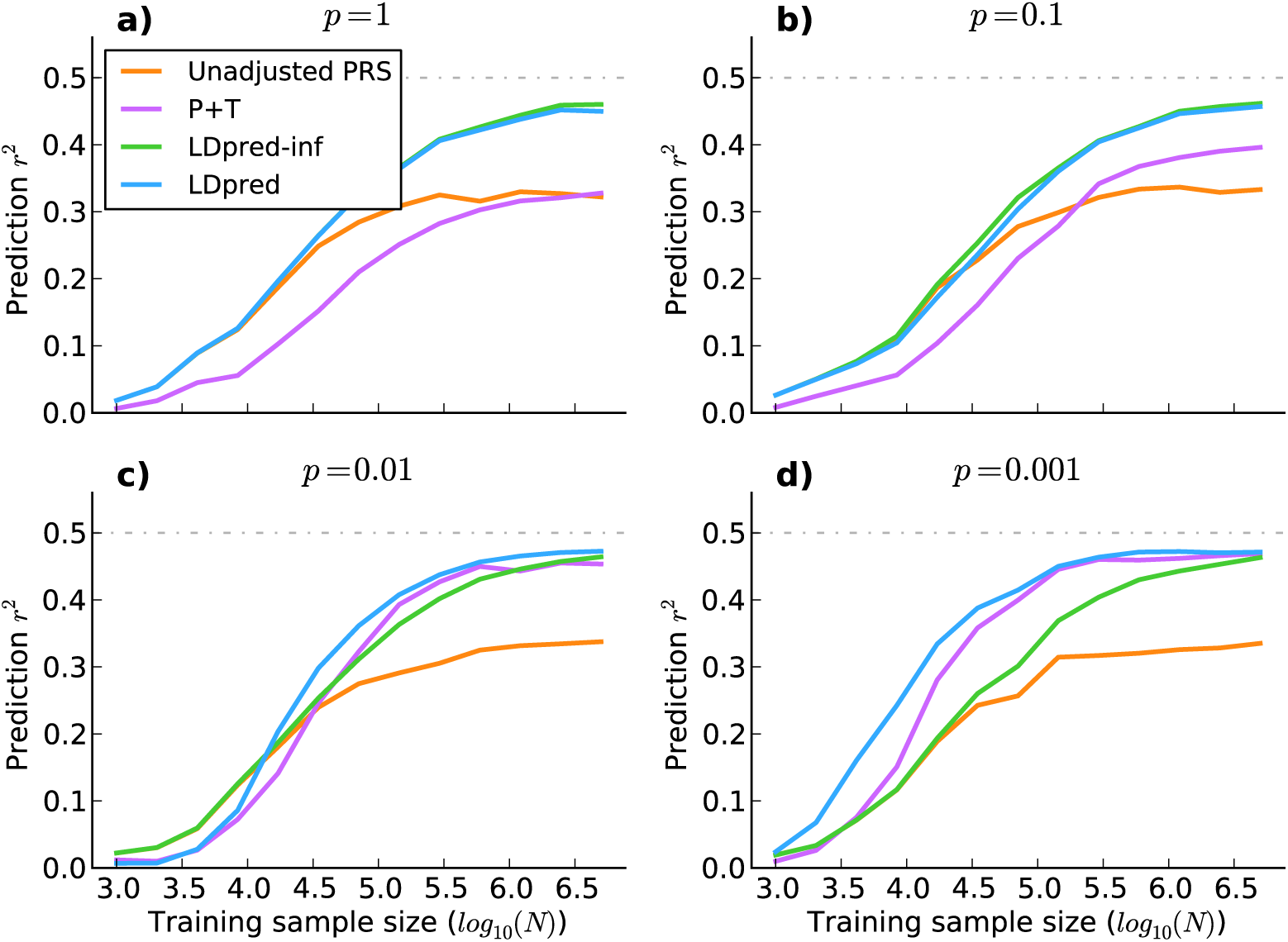
Comparison of methods using simulated genotypes (see **Supplementary Figure 4.**) with LD with Laplace mixture distributed effects instead of Gaussian mixture distributed effects. The change in prior appears to have minimal effect on the shape of the curve and the relative performance.

**Supplementary Figure 6.**
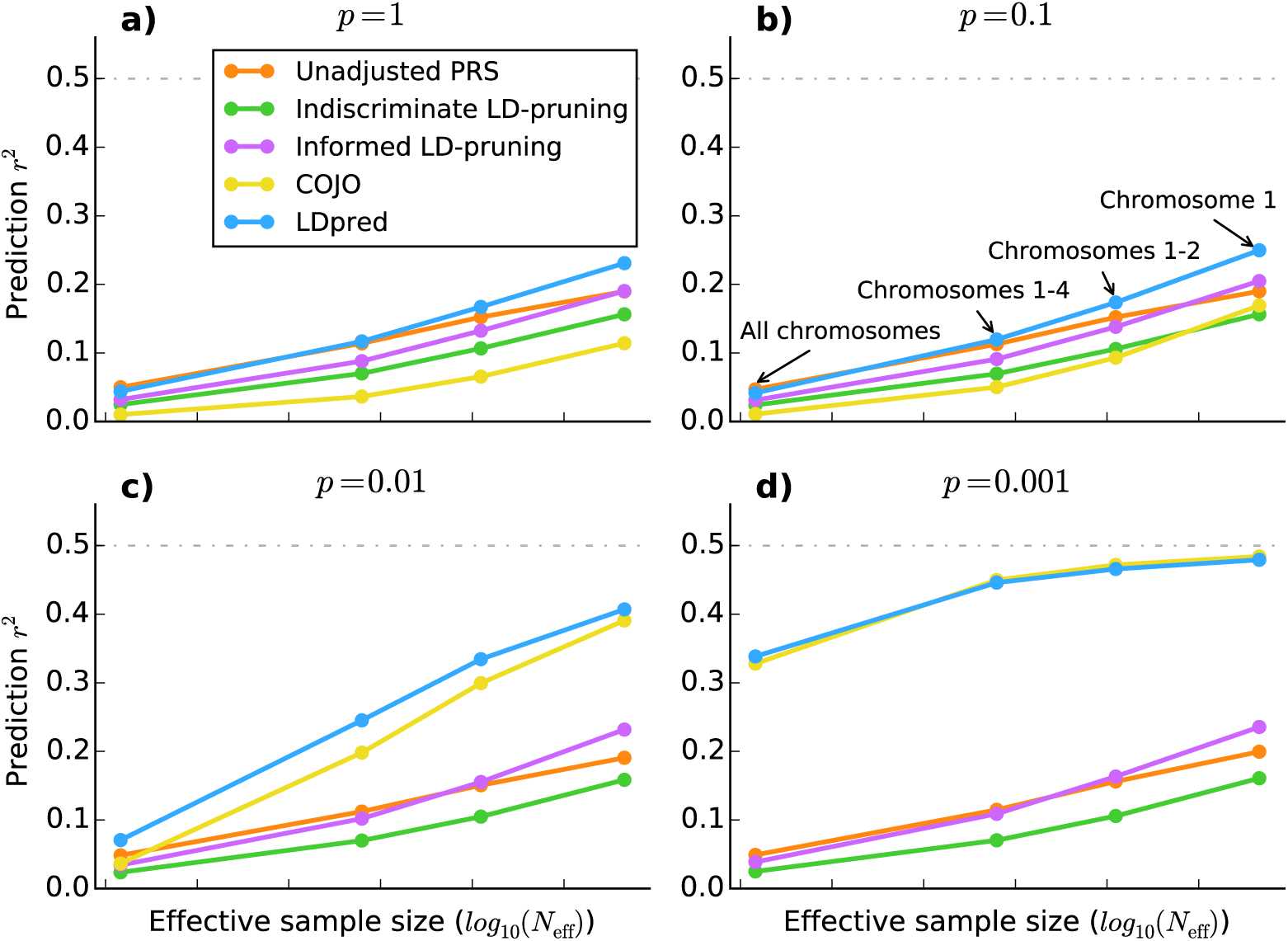
Comparisons to other methods using simulated traits and real WTCCC genotypes. As expected COJO^14,35^ performs close to optimal with sufficient training data, or more precisely, when the ratio *(Nh^2^)/(Mp)* is approximately larger than 10. The comparison between the two types of LD-pruning clearly demonstrates the advantage of informed LD-pruning over indiscriminate LD-pruning, which randomly prunes either marker of a pair of markers in LD. For both LD-pruning strategies a pair of markers was considered in LD if *r^2^*>0.2. When LDpred is compared to conditional joint analysis (COJO), LDpred outperforms COJO as long as the data does not overwhelm the prior, i.e. when *(Nh^2^)/(Mp)* is not sufficiently large (<10). For most of the diseases considered in this paper, current sample sizes are still not large enough for joint estimates to yield accurate risk scores.

**Supplementary Figure 7.**
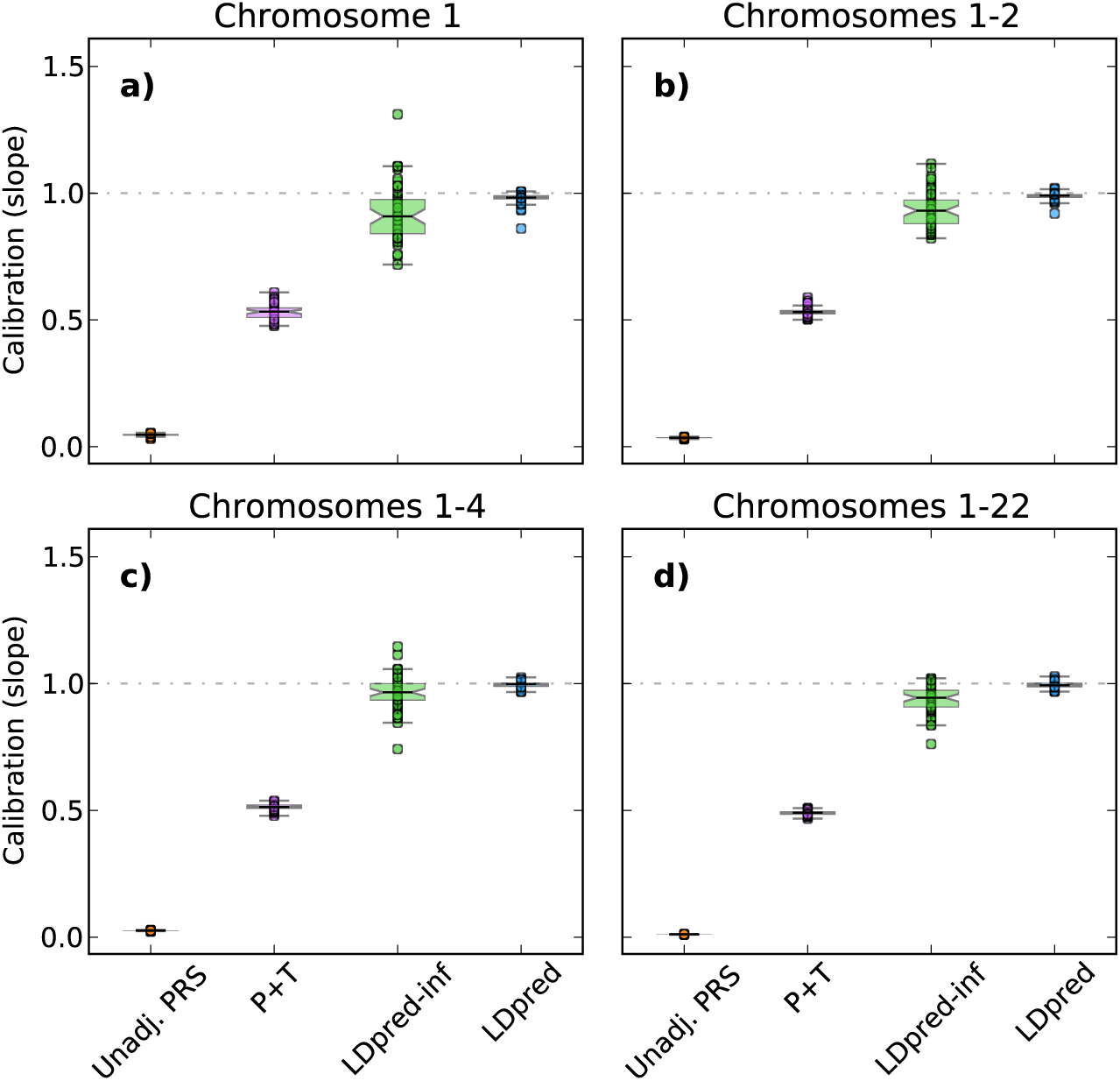
Boxplots of calibration slopes for the four prediction methods evaluated in Figure 2 for *p=0.001* (the fraction of variants with non-zero effects). The subfigures **a-d** correspond to different number of SNPs used, ranging from 30,004 SNPs on chromosome 1 in **a)** to 376,901 SNPs or the full genome in **d)**. If the prediction conditional on the true value is unbiased then we expect a slope of one. A slope of less than one implies that the predicted value is mis-calibrated by a factor of 1/slope. Results for other values of *p(p=1; p=0.1; p=0.01)* gave similar results, and even stronger bias for P+T (LD-pruning followed by *P*-value thresholding).

**Supplementary Table 1.** Numerical values of results displayed in Figure 3, on four different *R^2^* or AUC scales. To transform the prediction *R^2^* to liability scale we used the Lee *et al. R^2^* transformation^36^ using values of disease prevalence specified in Supplementary Table 2.

| Disease | Prediction accuracy measurement | Unadjusted PRS using all SNPs | Pruning + Thresholding | LDpred-inf | LDpred |
| --- | --- | --- | --- | --- | --- |
| **T1D** | Observed scale $R^2$ | 0.1064 | 0.3195 | 0.1062 | 0.3832 |
| | Nagelkerke $R^2$ | 0.1442 | 0.4228 | 0.1438 | 0.5084 |
| | Liability scale $R^2$ | 0.0426 | 0.0934 | 0.0426 | 0.1037 |
|  | AUC | 0.6915 | 0.8410 | 0.6912 | 0.8738 |
| **T2D** | Observed scale $R^2$ | 0.0360 | 0.0465 | 0.0404 | 0.0467 |
| | Nagelkerke $R^2$ | 0.0488 | 0.0631 | 0.0547 | 0.0633 |
| | Liability scale $R^2$ | 0.0257 | 0.0327 | 0.0287 | 0.0329 |
|  | AUC | 0.6094 | 0.6243 | 0.6180 | 0.6275 |
| **CAD** | Observed scale $R^2$ | 0.0250 | 0.0349 | 0.0290 | 0.0333 |
| | Nagelkerke $R^2$ | 0.0338 | 0.0473 | 0.0393 | 0.0451 |
| | Liability scale $R^2$ | 0.0191 | 0.0263 | 0.0221 | 0.0253 |
|  | AUC | 0.5880 | 0.6087 | 0.5963 | 0.6043 |
| **CD** | Observed scale $R^2$ | 0.0428 | 0.0485 | 0.0461 | 0.0824 |
| | Nagelkerke $R^2$ | 0.0585 | 0.0661 | 0.0630 | 0.1122 |
| | Liability scale $R^2$ | 0.0148 | 0.0167 | 0.0159 | 0.0267 |
|  | AUC | 0.6212 | 0.6313 | 0.6279 | 0.6693 |
| **RA** | Observed scale $R^2$ | 0.0483 | 0.1151 | 0.0462 | 0.1354 |
| | Nagelkerke $R^2$ | 0.0656 | 0.1540 | 0.0627 | 0.1801 |
| | Liability scale $R^2$ | 0.0239 | 0.0508 | 0.0229 | 0.0579 |
|  | AUC | 0.6277 | 0.6994 | 0.6267 | 0.7162 |
| **BD** | Observed scale $R^2$ | 0.0707 | 0.0876 | 0.0798 | 0.0816 |
| | Nagelkerke $R^2$ | 0.09578 | 0.1185 | 0.1080 | 0.1105 |
| | Liability scale $R^2$ | 0.0308 | 0.0371 | 0.0342 | 0.0349 |
|  | AUC | 0.6552 | 0.6744 | 0.6662 | 0.6682 |
| **HT** | Observed scale $R^2$ | 0.0306 | 0.0424 | 0.0348 | 0.0376 |
| | Nagelkerke $R^2$ | 0.0414 | 0.0574 | 0.0471 | 0.0509 |
| | Liability scale $R^2$ | 0.0258 | 0.0351 | 0.0292 | 0.0314 |
|  | AUC | 0.6005 | 0.6180 | 0.6072 | 0.6109 |

**Supplementary Table 2.** P+T and LDpred parameters for methods evaluated in Figure 3. The heritabilities are calculated as averages over 5 cross validations. The Lee *et al*. heritability transformation^52^ was used to obtain the heritability on the liability scale. The LD window size used in the simulations was 400 SNPs.

| Disease | Optimal fraction of causal markers used in LDpred | Optimal *P*-value threshold for Pruning + Thresholding | LDpred estimated heritability | LDpred estimated heritability on liability scale | Assumed disease prevalence |
| --- | --- | --- | --- | --- | --- |
| **T1D** | 0.001 | 10<sup>-6</sup> | 1.3250 | 0.7258 | 0.005 |
| **T2D** | 0.03 | 1 | 0.6206 | 0.5125 | 0.03 |
| **CAD** | 0.03 | 1 | 0.6160 | 0.5181 | 0.035 |
| **CD** | 0.01 | 0.0001 | 0.7974 | 0.2904 | 0.001 |
| **RA** | 0.0001 | 10<sup>-6</sup> | 0.9097 | 0.5145 | 0.0075 |
| **BD** | 0.1 | 1 | 0.9695 | 0.4959 | 0.005 |
| **HT** | 0.03 | 1 | 0.6216 | 0.5939 | 0.05 |

**Supplementary Table 3.**
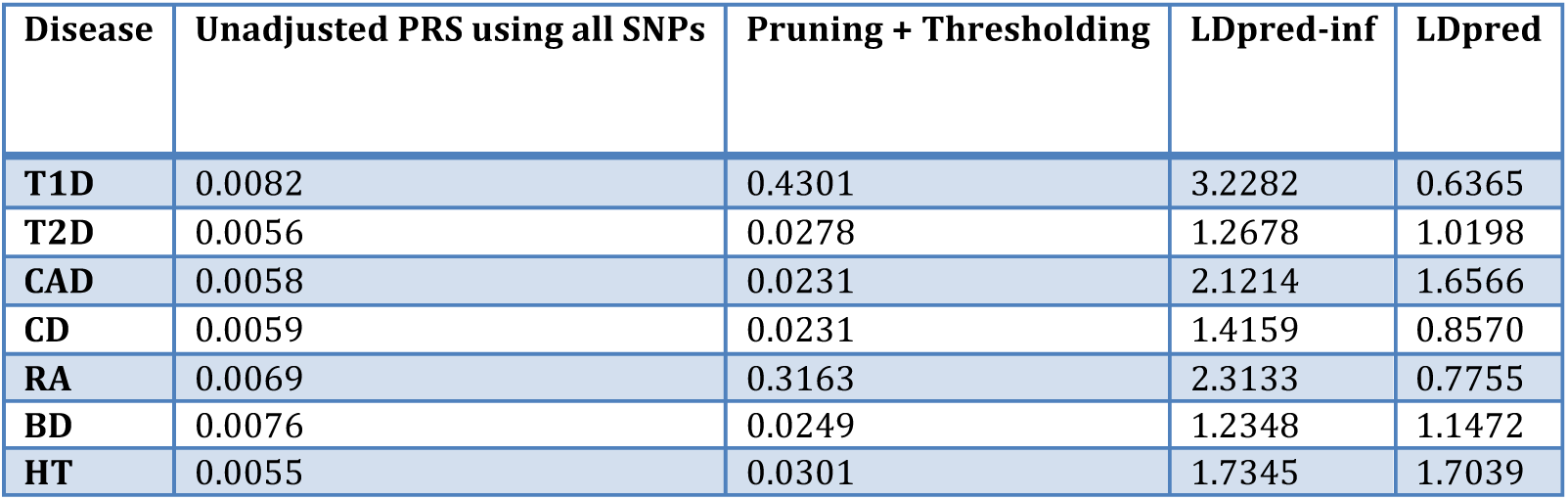
Calibration comparison for methods evaluated in Figure 3. We report the slope, where a value close to 1 represents a well-calibrated prediction. LDpred yields the most appropriately calibrated predictions.

| Disease | Unadjusted PRS using all SNPs | Pruning + Thresholding | LDpred-inf | LDpred |
| --- | --- | --- | --- | --- |
| **T1D** | 0.0082 | 0.4301 | 3.2282 | 0.6365 |
| **T2D** | 0.0056 | 0.0278 | 1.2678 | 1.0198 |
| **CAD** | 0.0058 | 0.0231 | 2.1214 | 1.6566 |
| **CD** | 0.0059 | 0.0231 | 1.4159 | 0.8570 |
| **RA** | 0.0069 | 0.3163 | 2.3133 | 0.7755 |
| **BD** | 0.0076 | 0.0249 | 1.2348 | 1.1472 |
| **HT** | 0.0055 | 0.0301 | 1.7345 | 1.7039 |

**Supplementary Table 4.** Numerical values of results displayed in Figure 4, on four different *R*^2^ or AUC scales.

| Disease | Prediction accuracy measurement | Unadjusted PRS using all SNPs | Pruning + Thresholding | LDpred-inf | LDpred |
| --- | --- | --- | --- | --- | --- |
| **SCZ-MGS** | Observed scale $R^2$ | 0.1591 | 0.1510 | 0.1870 | 0.1898 |
| | Nagelkerke $R^2$ | 0.2119 | 0.2014 | 0.2488 | 0.2528 |
| | Liability scale $R^2$ | 0.0616 | 0.0594 | 0.0688 | 0.0694 |
|  | AUC | 0.7294 | 0.7248 | 0.7499 | 0.7519 |
| **SCZ-ISC** | Observed scale $R^2$ | 0.1169 | 0.0970 | 0.1334 | 0.1367 |
| | Nagelkerke $R^2$ | 0.1574 | 0.1304 | 0.1803 | 0.1836 |
| | Liability scale $R^2$ | 0.0518 | 0.0446 | 0.0578 | 0.0585 |
|  | AUC | 0.6988 | 0.6784 | 0.7127 | 0.7165 |
| **MS** | Observed scale $R^2$ | 0.0316 | 0.0674 | 0.0363 | 0.0840 |
| | Nagelkerke $R^2$ | 0.0474 | 0.0978 | 0.0512 | 0.1198 |
| | Liability scale $R^2$ | 0.0149 | 0.0302 | 0.0170 | 0.0368 |
|  | AUC | 0.6169 | 0.6714 | 0.6187 | 0.6918 |
| **BC** | Observed scale $R^2$ | 0.0071 | 0.0324 | 0.0092 | 0.0386 |
| | Nagelkerke $R^2$ | 0.0097 | 0.0437 | 0.0119 | 0.0519 |
| | Liability scale $R^2$ | 0.0040 | 0.0184 | 0.0052 | 0.0220 |
|  | AUC | 0.5489 | 0.6052 | 0.5549 | 0.6156 |
| **T2D** | Observed scale $R^2$ | 0.0159 | 0.0247 | 0.0214 | 0.0273 |
| | Nagelkerke $R^2$ | 0.0212 | 0.0330 | 0.0309 | 0.0365 |
| | Liability scale $R^2$ | 0.0112 | 0.0170 | 0.0149 | 0.0187 |
|  | AUC | 0.5747 | 0.5854 | 0.5825 | 0.5953 |
| **CAD** | Observed scale $R^2$ | 0.0109 | 0.0101 | 0.0124 | 0.0125 |
| | Nagelkerke $R^2$ | 0.0146 | 0.0137 | 0.0168 | 0.0170 |
| | Liability scale $R^2$ | 0.0085 | 0.0080 | 0.0097 | 0.0098 |
|  | AUC | 0.5612 | 0.5557 | 0.5645 | 0.5647 |

**Supplementary Table 5.** Parameters inferred or assumed by P+T and LDpred for results displayed in Figure 4. The Lee *et al*. heritability transformation^52^ was used to obtain the heritablity on the liability scale.

| Disease | Optimal $P$-value threshold for Pruning + Thresholding | Optimal Gaussian mixture weight (fraction of causal markers) for LDpred | LDpred/LD-pruning window size (# of SNPs) | GWAS sample size used in LDpred | LDpred estimated heritability | LDpred estimated heritability on liability scale | Assumed prevalence |
| --- | --- | --- | --- | --- | --- | --- | --- |
| SCZ-MGS | 0.1 | 0.3 | 500 | 65K | 0.5738 | 0.4231 | 0.01 |
| SCZ-ISC | 0.1 | 0.3 | 500 | 65K | 0.4718 | 0.3479 | 0.01 |
| MS | 0.001 | 0.01 | 400 | 27K | 0.3694 | 0.1321 | 0.001 |
| BC | 0.00003 | 0.003 | 400 | 50K | 0.1934 | 0.1124 | 0.01 |
| T2D | 0.00003 | 0.1 | 300 | 69K | 0.2061 | 0.1582 | 0.0075 |
| CAD | 0.1 | 1 | 300 | 86K | 0.2943 | 0.2494 | 0.035 |

**Supplementary Table 6.** Calibration slopes for methods evaluated in Figure 4. We report the slope, where a value close to 1 represents a well-calibrated prediction.

| Disease | Unadjusted PRS using all SNPs | Pruning + Thresholding | LDpred-inf | LDpred |
| --- | --- | --- | --- | --- |
| SCZ-MGS | 0.0063 | 0.0467 | 0.3845 | 0.3918 |
| SCZ-ISC | 0.0130 | 0.0407 | 0.4683 | 0.4413 |
| MS | 0.0089 | 0.0717 | 0.9092 | 0.2011 |
| BC | 0.0017 | 0.1327 | 1.2323 | 0.5650 |
| T2D | 0.0032 | 0.1002 | 0.6421 | 0.4057 |
| CAD | 0.0035 | 0.0137 | 0.2244 | 0.1868 |

**Supplementary Table 7.** Numerical values of results displayed in Figure 4, on four different *R*^2^ or AUC scales.

| Schizophrenia Cohort | Prediction accuracy measurement | Unadjusted PRS using all SNPs | Pruning + Thresholding | LDpred-inf | LDpred |
| --- | --- | --- | --- | --- | --- |
| **MGS (European ancestry)** | Observed scale $R^2$ | 0.1591 | 0.1510 | 0.1870 | 0.1898 |
| | Nagelkerke $R^2$ | 0.2119 | 0.2014 | 0.2488 | 0.2528 |
| | Liability scale $R^2$ | 0.0616 | 0.0594 | 0.0688 | 0.0694 |
|  | AUC | 0.7294 | 0.7248 | 0.7499 | 0.7519 |
| **JPN1 (Japanese ancestry)** | Observed scale $R^2$ | 0.0477 | 0.0702 | 0.0691 | 0.0695 |
| | Nagelkerke $R^2$ | 0.0635 | 0.0944 | 0.0923 | 0.0929 |
| | Liability scale $R^2$ | 0.0232 | 0.0323 | 0.0319 | 0.0320 |
|  | AUC | 0.6276 | 0.6527 | 0.6523 | 0.6531 |
| **TCR1 (Chinese ancestry)** | Observed scale $R^2$ | 0.0570 | 0.0616 | 0.0704 | 0.0717 |
| | Nagelkerke $R^2$ | 0.0761 | 0.0821 | 0.0939 | 0.0956 |
| | Liability scale $R^2$ | 0.0274 | 0.0294 | 0.0329 | 0.0336 |
|  | AUC | 0.6331 | 0.6391 | 0.6483 | 0.6488 |
| **HOK2 (Chinese ancestry)** | Observed scale $R^2$ | 0.0253 | 0.0306 | 0.0374 | 0.0373 |
| | Nagelkerke $R^2$ | 0.0414 | 0.0511 | 0.0609 | 0.0609 |
| | Liability scale $R^2$ | 0.0187 | 0.0225 | 0.0271 | 0.0271 |
|  | AUC | 0.6176 | 0.6250 | 0.6352 | 0.6352 |
| **AFAM (African American ancestry)** | Observed scale $R^2$ | 0.0170 | 0.0151 | 0.0279 | 0.0280 |
| | Nagelkerke $R^2$ | 0.0233 | 0.0202 | 0.0382 | 0.0383 |
| | Liability scale $R^2$ | 0.0095 | 0.0084 | 0.0152 | 0.0152 |
|  | AUC | 0.5745 | 0.5682 | 0.5936 | 0.5936 |

**Supplementary Table 8.**
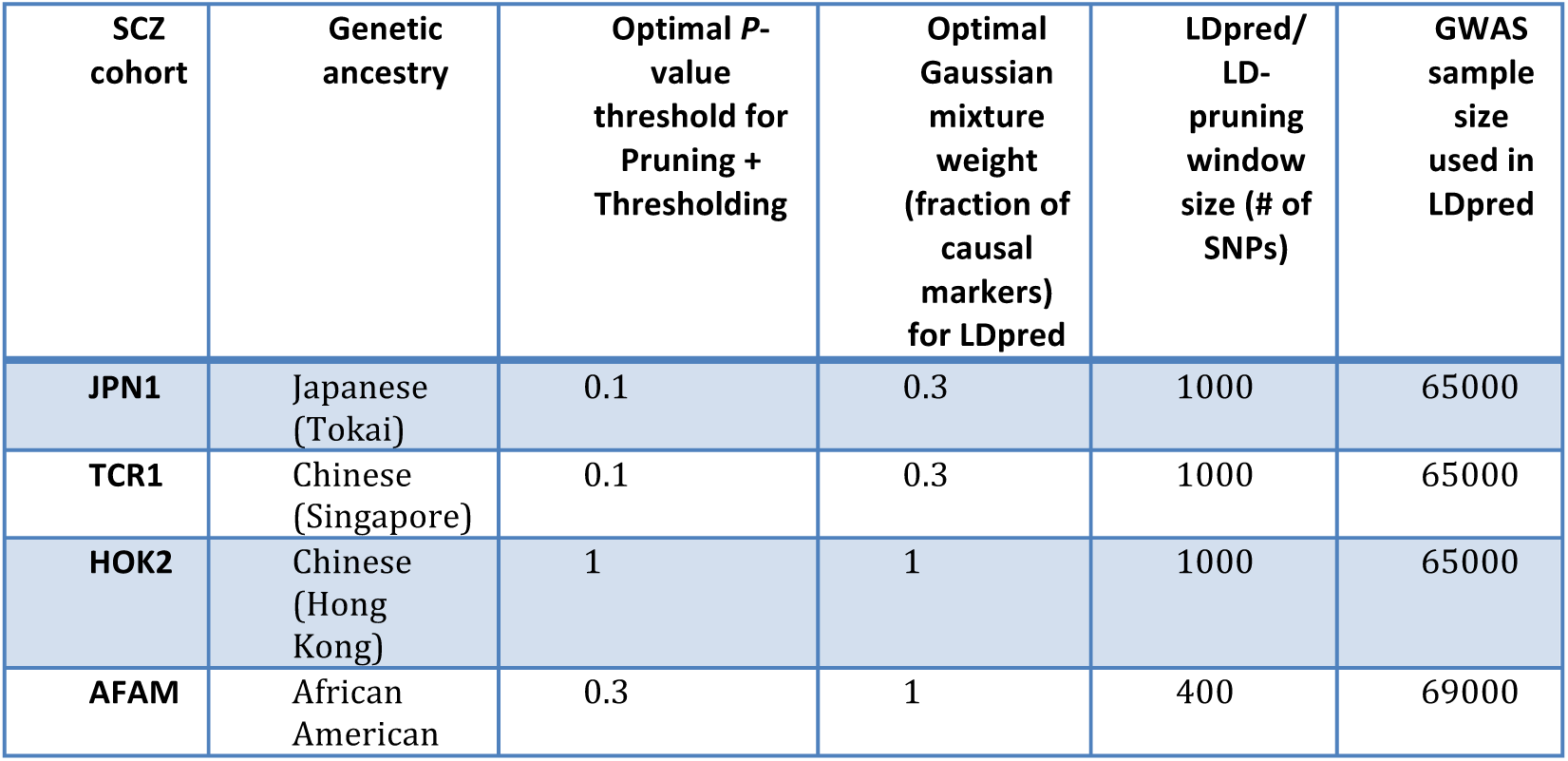
Parameters inferred or assumed by P+T and LDpred for analysis of the non-European validation samples in **Supplementary Table 7**.

## 1 Phenotypic model and notation

Throughout this paper we model the phenotypes as additive unless specified otherwise. This is an important simplifying assumption that is commonly made in the statistical genetics literature, e.g. when estimating heritability with variance components [1]. Let *ϕ* denote a phenotype vector with *N* values. Then the phenotype can be written as follows

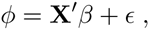

where **X** denotes a *M* × *N* genotype matrix with *M* genotypes values for *N* individuals, *β* is a *M* vector of genotype effects, and *ϵ* is the residual noise. For simplicity, we assume that the genotypes and the phenotype have been normalized appropriately, i.e. E(*ϕ*) = 0, Var(*ϕ*) = 1 and ∀*_i_* E(**X***_i_*) = 0, Var(**X***_i_*) = 1. Finally, assume that *ϵ* is Gaussian distributed *ϵ* ~ *N*(0, (1 − *h*^2^)**I**), where *h*^2^ is the heritability of the trait. Under the liability threshold model [2], binary traits, such as case-control phenotypes can be modeled as quantitative by using a transformation to the liability scale.

## 2 Posterior mean phenotype estimation (LDpred and LDpred-inf)

Under the assumption that the phenotype has an additive genetic architecture and is linear, then estimating the posterior mean phenotype boils down to estimating the posterior mean effects of each SNP and then summing their contribution up in a risk score.

### 2.1 Posterior mean effects assuming unlinked markers and an infinitesimal model

We will first consider the infinitesimal model, which represents a genetic architecture where all genetic variants are causal. The classical example is Fisher’s infinitesimal model [3], which assumes genotypes are unlinked effect sizes have a Gaussian distribution (after normalizing by allele frequency).

#### Gaussian prior (infinitesimal model)

Assume that *β_i_* are independently drawn from a Gaussian distribution 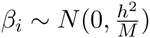, where *M* denotes the total number of causal effects (*β_i_*), and *N* the number of individuals. Then we can derive a posterior mean given the ordinary least square estimate 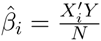. The least square estimate is approximately distributed as
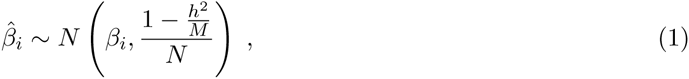

where the variance can be approximated further, 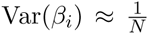, when *M* is large. Using this the posterior distribution for *β_i_* is
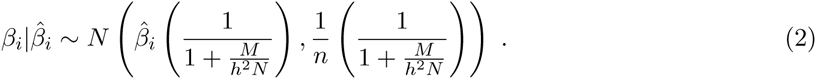

This suggest that a uniform shrink by a factor of 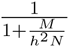 is appropriate under Fisher’s infinitesimal model.

#### Laplace prior (infinitesimal model)

Under the Fisher/Orr model, causal effects are expected to be approximately exponentially distributed [4]. Empirical evidence largely supports this for human diseases, but also points to a genetic architecture in which there are fewer large effects [5]. Regardless, a double Exponential or a Laplace distribution is arguably a reasonable prior distribution for the effect sizes, where the variance is 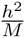 (so that they sum up to the total heritability). Under this model, the probability density function for *β_i_* becomes
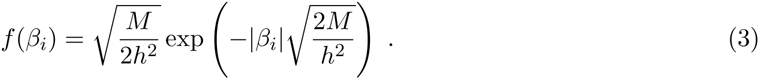

Using the Bayes theorem we can write out the posterior density given the ordinary least square estimate as follows
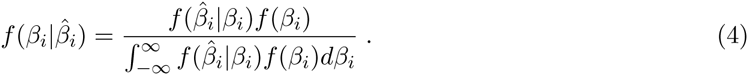

Using the fact that the ordinary least square estimates are Gaussian distributed as shown in equation 2.1, we can write out the term in the integral as follows
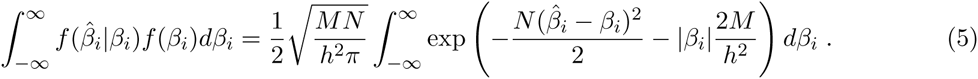

This integral is non-trivial, however we can solve it numerically [6]. Similarly, the posterior mean, 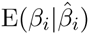, also yields a non-trivial integral that can be evaluated numerically.

#### LASSO shrink

When the effects have a Gaussian prior distribution the posterior prior is symmetric, causing mean and mode to be equal. This is not the case when we use a Laplace prior for the effects. Although the posterior mean requires numerical integration, it turns out that the posterior mode has a simple analytical form [7]. The posterior mode under a Laplace prior is in fact the LASSO estimate [8]. If we assume that the sum of the effects has variance *h*^2^, and that the genetic markers are uncorrelated, then the posterior mode estimate is
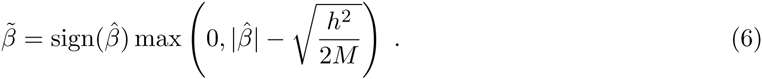

Interestingly, the posterior mode effects for estimated effects below a given threshold are set to 0, even though all betas are causal in the model.

**Figure 1:**
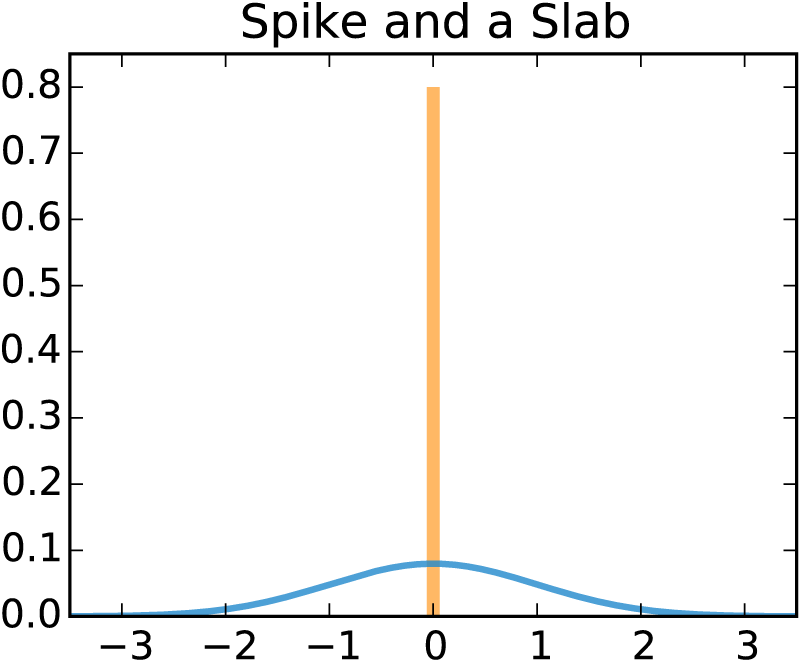
An illustration of a spike and slab prior with a Gaussian slab.

### 2.2 Posterior mean effects assuming unlinked markers and a non-infinitesimal model

Most diseases and traits are not likely to be strictly infinitesimal, i.e. follow Fisher’s infinitesimal model [3]. Instead, a non-infinitesimal model, where only a fraction of the genetic variants are truly causal and affect the trait, is more likely to describe the underlying genetic architecture. We can model non-infinitesimal genetic architectures using mixture distributions with a mixture parameter *p* that denotes the fraction of causal markers. More specifically, we will consider a spike and slab prior with a 0-spike and Gaussian slab.

#### Gaussian mixture prior (spike and a slab)

Assume that the effects are drawn from a mixture distribution as follows.
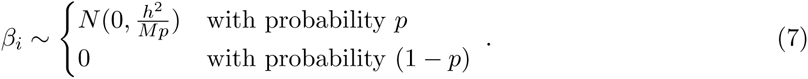

Another way of writing this is to use Dirac’s delta function, i.e. write *β_i_* = *pu* + (1 − *p*)*v*, where 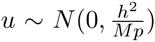 and 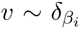. Here 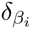 denotes the point density at *β_i_* = 0, which integrates to 1. We can then write out the density for 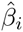 as follows
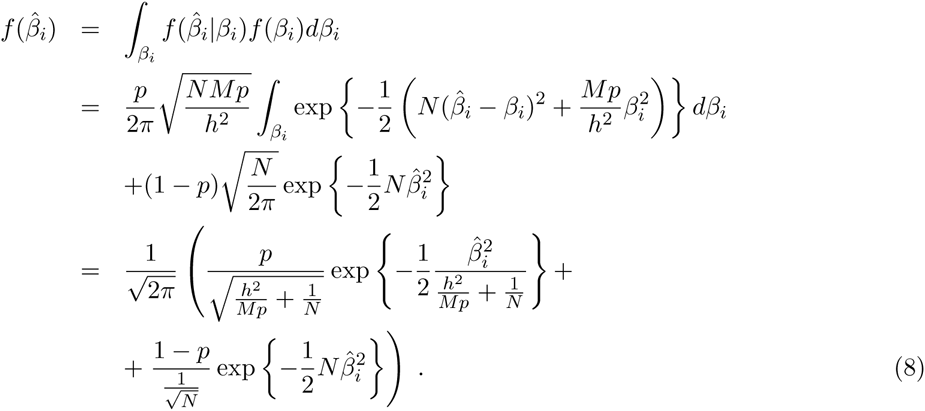

We are interested in the posterior mean which can be expressed as
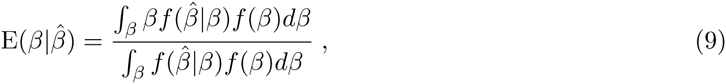

hence we only need to calculate the following definite integral
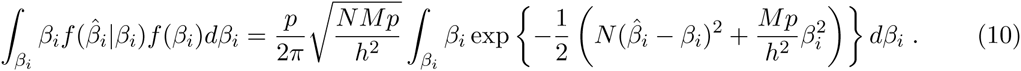

Thus the posterior mean is
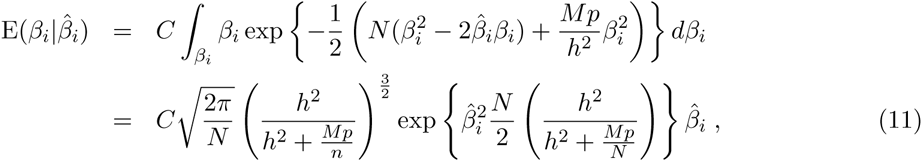

where
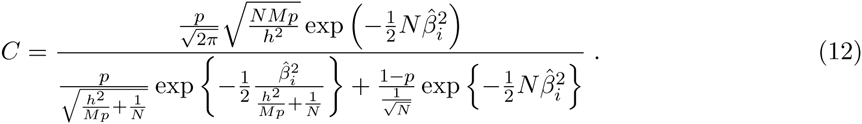

Now by realizing that the posterior probability that *β_i_* is sampled from the Gaussian distribution given 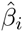 is exactly
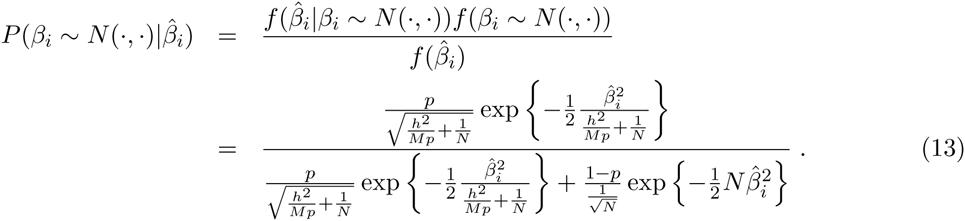

then we can rewrite the posterior mean in a simpler fashion. If we let 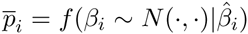, denote the posterior probability that *β_i_* is non-zero or Gaussian distributed, then it becomes
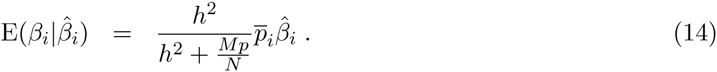

### 2.3 Posterior mean effects assuming linked markers and an infinitesimal model (LDpred-inf)

Following Yang *et al*. [9], we have can obtain the joint least square effect estimates as
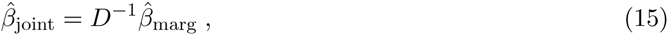

where 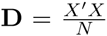 is the LD matrix. In practice, the LD matrix is *M* × *M* and likely singular, e.g. if two (or more) markers are in perfect linkage. If the LD matrix **D** is singular, there the joint least square estimate does not have a unique solution. However, if the individuals in the training data do not display family or population structure, the genome-wide LD matrix is approximately a banded matrix, which allows adjust for LD locally instead. To formalize these ideas, let us introduce some notation. Let *l_i_* denote the *i*’th locus or region with 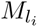 markers, and let 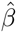 denote the marginal least square estimate vector unless otherwise noted. In addition, let *β*^(*i*)^ denote the vector of effects that are in the *i*’th region, and similarly 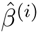 the corresponding vector of marginal effect estimates in the region. Under this model we can derive the posterior distribution for effect estimates at the *i*’th region, i.e. 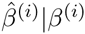. The posterior mean is 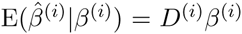, where 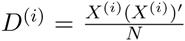 is the LD matrix obtained from the markers in the *i*’th region, i.e. *X*^(*i*)^. Futhermore, the conditional covariance matrix is
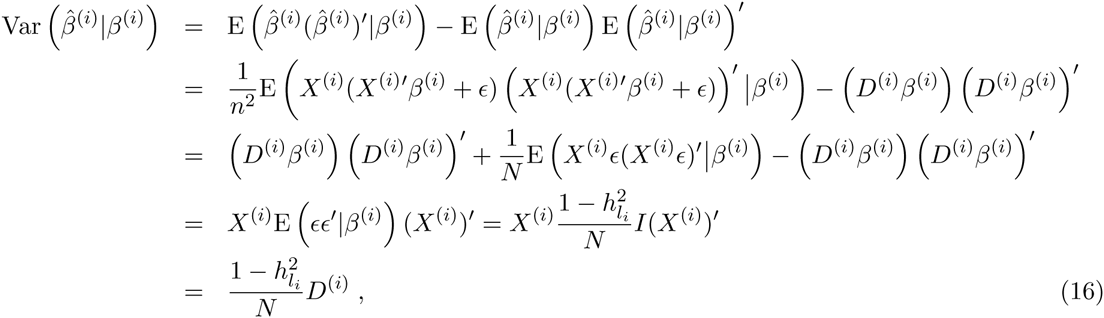

where 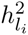 denotes the heritability explained by the markers in the region, i.e. *X*^(*i*)^. If we assume that the heritability explained by an individual region is small, then this simplifies to Var 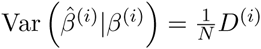. This equation is particularly useful for performing efficient simulations of effect sizes without simulating the genotypes. Given an LD matrix, *D*, we can simulate effect sizes and corresponding least square estimates. Similarly, for the joint estimate we have
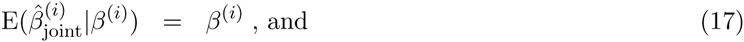

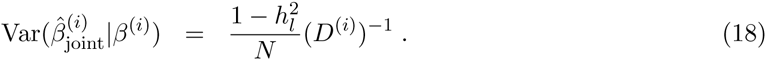

Since 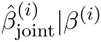 follows a multivariate Gaussian distribution, it follows that the maximum likelihood estimator of the jointly estimated marker effects given least square estimated marker effects (summary statistics) is 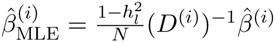.

#### Gaussian distributed effects

In the following, we let *β* (and respectively 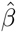) denote the effects within a region of LD. We furthermore assume that these markers only explain a fraction, 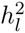, of the total phenotypic variance, and 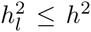. Given the prior distribution for *β* and the conditional distribution 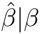 we can derive the posterior mean by considering the joint density:
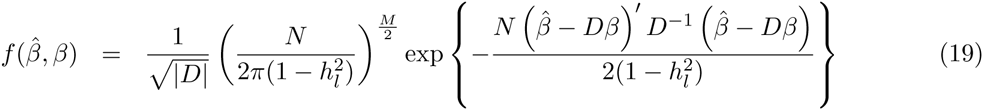

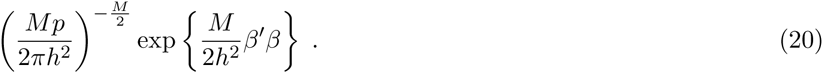

We can derive the posterior density for 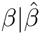 by completing the square in the exponential. This yields a multivariate Gaussian with mean and variance as follows
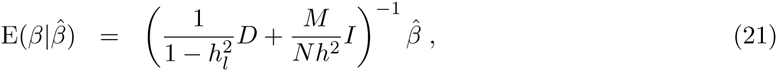

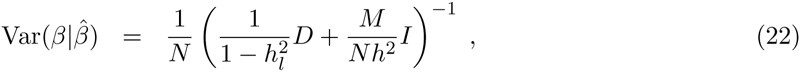

where *h*^2^ denotes the heritability explained by the *m* causal variants and 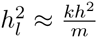 is the heritability of the *k* variants in *X*. If *M* ≫ *k*, then (1 − *h_l_*) becomes approximately one, and the equations above can be simplified accordingly. As expected, the posterior mean approaches the maximum likelihood estimator as the training sample size grows.

### 2.4 Posterior mean effects assuming linked markers and a non-infinitesimal model

In this section we will derive an LD-adjustment under a non-infinitesimal model.

#### Gaussian distributed effects

Define *q* as follows
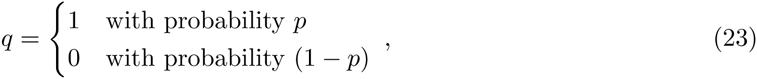

then we can write *β* = *qu*, where 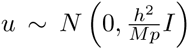. Hence we can write the multivariate density function for *β* as
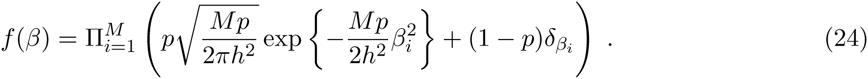

The posterior distribution for 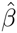 given *β* is
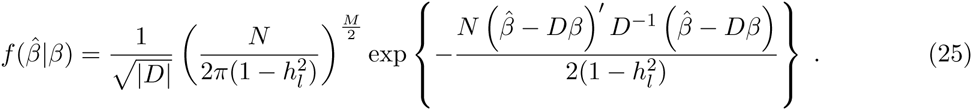

As usual, we want to calculate the posterior mean, i.e.
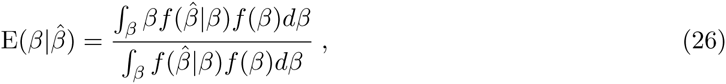

which consists of two *m* dimensional integrands. Any multiplicative term that does not involve *β* in the two integrands factors out. The integrands consist of 2*^M^* additive terms. We therefore result to a MCMC Gibbs sampler to sample from the posterior and estimate the posterior mean effects.

#### Metropolis Hastings Markov Chain Monte Carlo

An alternative approach to obtaining the posterior mean is to sample from the posterior distribution, and then average over the samples to obtain the posterior mean. In our case we know the posterior up to a constant, i.e.
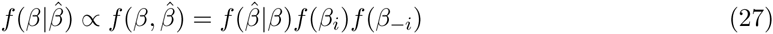

where *β_−i_* denotes all the other effects except for the effect of the *i*’th marker. We can use this fact to sample efficiently in a Markov chain Monte Carlo setting where we sample one marker effect at a time in an iterative fashion (the conditional proposal distribution is therefore univariate). This ensures that the Metropolis-Hastings acceptance ratio 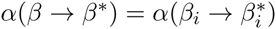 only depends on local LD, and not the distributions of other effects, i.e.
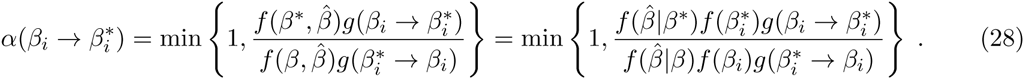

where the asterisk denotes the proposed effect as sampled from the conditional proposal distribution *g*. Since Driac’s delta density is infinite for a zero value, this ratio is undefined under the previously proposed infinitesimal model. Therefore, we consider an alternative mixture distribution with two Gaussians, one with variance (1 − *τ*)*h*^2^/(*mp*) and the other with variance *τh*^2^/(*m*(1 −*p*)) where *τ* is a small number, say *τ* = 10^−3^. Hence the prior distribution becomes
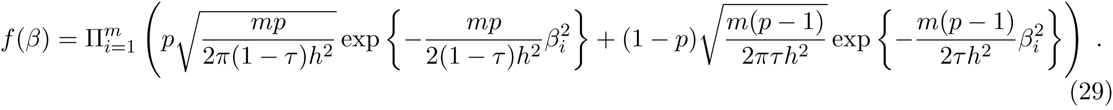

The conditional distribution 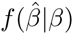 is still the same and is given in equation (25). Together this gives us all the quantities needed to implement the Metropolis Hastings MCMC.

#### Approximate Gibbs sampler (LDpred)

The general MH MCMC described above is tedious to implement and can also be computationally inefficient if proposal distributions are not carefully chosen. As a more efficient MCMC approach, we also considered a Gibbs sampler. This requires us to derive the marginal conditional posterior distributions for effects, i.e. 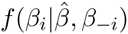, where *β_−i_* refers to the vector of betas excluding the *i*’th beta. We can write the posterior distribution as follows
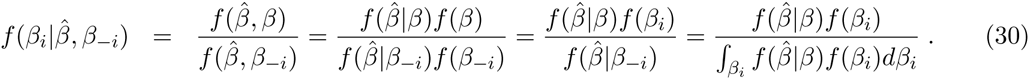

Sampling from this distribution is not trivial. However, we can partition the sampling procedure into two parts where we first sample whether the effect is different from 0 or not, and then if it is different from zero we can assume it has a Gaussian prior. To achieve this we first need to calculate the posterior probability of a marker being causal, i.e.
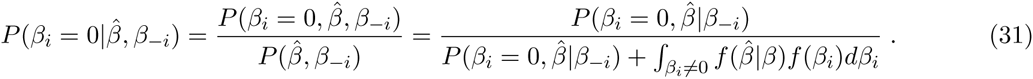

Obtaining an analytical solution to this is non-trivial, however, if we assume that 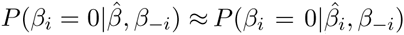, then we can simply extract out the effects of LD from other effects on the effect estimate 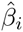 and then use the marginal posterior probability of the marker being causal from equation (13) instead, i.e. 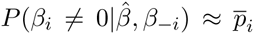. If we sample the effect to be non-zero, and again make the simplifying assumption that 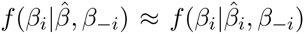 then its posterior distribution becomes simple and we can again extract the effects of LD on the effect estimate and sample from the marginal posterior distribution derived in section 2.2. To summarize the marginal posterior distribution for *β_i_* becomes
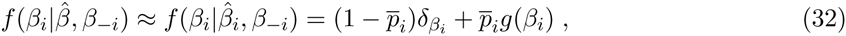

where *g*(*β_i_*) is the Gaussian density for the posterior distribution conditional on *β_i_* ≠ 0, i.e.
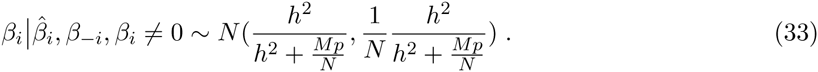

#### Estimating heritability from summary statistics

In the derivation above we assume that the heritability of the trait is known. Although one can usually find some heritability estimate in the literature for most traits, it is preferable to estimate it directly from the training data GWAS summary statistics. We achieve this by using LD score regression [?] to estimate the heritability explained by the genotyped SNPs in the GWAS summary statistics. The heritability estimate is
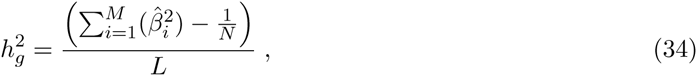

where 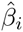 is the least square estimate for the *i*’th SNP (from the GWAS summary statistics) and *L* is the genome-wide average LD score. Further information on estimating heritability from GWAS summary statistics is provided in (Finucane *et al.*, in preparation). In simulations, we found LDpred to be rather robust to minor changes in the heritability estimate.

#### Practical considerations for LDpred

Throughout the derivation of LDpred above we assumed that the LD information in the training data was known. However, in practice that information may not be available and instead we need to estimate the LD pattern from a reference panel. In simulations we found that the accuracy of this estimation does affect the performance of LDpred, and we recommend that the LD be estimated from reference panels with at least 2000 individuals. In the current implementation of LDpred we fixed an LD window around the genetic variant when calculating the posterior mean effect. This is a parameter in the model which the user can set, and the optimal value may depend on the number of markers and other factors. For our analysis we accounted for LD between the SNP and a fixed window of SNPs of each side. The actual number of SNPs that were used to account for LD depends on the total number of SNPs used in each analysis, with larger windows for larger datasets.

Although LDpred aims to estimate the posterior mean phenotype (the best unbiased prediction) it is only guaranteed to do so if all the assumptions hold. As LDpred relies on a few assumptions (both regarding LD and mathematical approximations), it is an *approximate* Gibbs sampler which can lead to robustness issues. Indeed, we found the LDpred to be sensitive to inaccurate LD estimates, especially for very large sample sizes. To address this we set the the probability of setting the effect size to 0 in the Markov chain to be at least 5%. This lead to a robust version of LDpred which performed well on both simulations and on real data. If converge issues arise when applying LDpred to other data, then it may be worthwhile to explore higher values for the 0-jump probability.

Finally, an important parameter that LDpred assumes to known is *p*, the fraction of “causal markers”. This parameter may of course not actually reflect the true fraction of causal markers as the model assumptions are, as always, flawed and the causal markers may not necessarily be genotyped. However, it is likely related to the true number of causal sites and may give valuable insight into the genetic architecture. Analogous to P-value thresholding we recommend that users calculate generate multiple LDpred polygenic risk scores for different values of *p* and then inferring and/or optimize on it in an independent validation data.

## 3 Conditional joint analysis

To understand the conditional joint (COJO) analysis as proposed by Yang et al. [1], we implemented a stepwise conditional joint analysis method in LDpred. The COJO analysis estimates the joint least square estimate from the marginal least square estimate (obtained from GWAS summary statistics). If we define 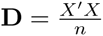, then we have the following relationship
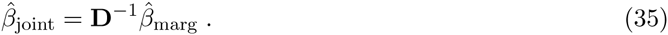

This matrix **D** has dimensions *M* × *M* and is likely to be singular in practice. However, as we do for LDpred, we can adjust for LD locally if the individuals in the training data do not display family or population structure, in which case the genome-wide LD matrix is approximately a banded matrix. In practice, the COJO analysis with all SNPs suffers a fundamental problem of statistical inference, i.e. it infers a large number of parameters (*M*) using *N* samples. Hence, if *N* < *M*, we do not expect the method to perform particularly well. We verified this using simulated effect estimates and validation genotypes (data not shown). By restricting to“top” SNPs and accounting for LD using a stepwise approach (as proposed by [1]) we alleviate this concern. However, although this reduces the overfitting problem when *N* < *M* this approach also risks discarding potentially informative markers from the analysis. Nevertheless, by optimizing the stopping threshold via cross-validation in an independent dataset, the method performs reasonably well in practice, especially when the number of causal markers in the genome is small. In contrast, LDpred conditions on the sample size and accounts for the noise term appropriately (under the model), leading to improved prediction accuracies regardless of training sample size.

## 4 Members of the Schizophrenia Working Group of the Psychiatric Genetics Consortium

The members of the Schizophrenia Working Group of the Psychiatric Genetics Consortium are Stephan Ripke, Benjamin M. Neale, Aiden Corvin, James T.R. Walters, Kai-How Farh, Peter A. Holmans, Phil Lee, Brendan Bulik-Sullivan, David A. Collier, Hailiang Huang, Tune H. Pers, Ingrid Agartz, Esben Agerbo, Margot Albus, Madeline Alexander, Farooq Amin, Silviu A. Bacanu, Martin Begemann, Richard A. Belliveau, Jr., Judit Bene, Sarah E. Bergen, Elizabeth Bevilacqua, Tim B. Bigdeli, Donald W. Black, Richard Bruggeman, Nancy G. Buccola, Randy L. Buckner, William Byerley, Wiepke Cahn, Guiqing Cai, Dominique Campion, Rita M. Cantor, Vaughan J. Carr, Noa Carrera, Stanley V. Catts, Kimberly D. Chambert, Raymond C.K. Chan, Ronald Y.L. Chen, Eric Y.H. Chen, Wei Cheng, Eric F.C. Cheung, Siow Ann Chong, C. Robert Cloninger, David Cohen, Nadine Cohen, Paul Cormican, Nick Craddock, James J. Crowley, David Curtis, Michael Davidson, Kenneth L. Davis, Franziska Degenhardt, Jurgen Del Favero, Lynn E. DeLisi, Ditte Demontis, Dimitris Dikeos, Timothy Dinan, Srdjan Djurovic, Gary Donohoe, Elodie Drapeau, Jubao Duan, Frank Dudbridge, Naser Durmishi, Peter Eichhammer, Johan Eriksson, Valentina Escott-Price, Laurent Essioux, Ayman H. Fanous, Martilias S. Farrell, Josef Frank, Lude Franke, Robert Freedman, Nelson B. Freimer, Marion Friedl, Joseph I. Friedman, Menachem Fromer, Giulio Genovese, Lyudmila Georgieva, Elliot S. Gershon, Ina Giegling, Paola Giusti-Rodrguez, Stephanie Godard, Jacqueline I. Goldstein, Vera Golimbet, Srihari Gopal, Jacob Gratten, Jakob Grove, Lieuwe de Haan, Christian Hammer, Marian L. Hamshere, Mark Hansen, Thomas Hansen, Vahram Haroutunian, Annette M. Hartmann, Frans A. Henskens, Stefan Herms, Joel N. Hirschhorn, Per Hoffmann, Andrea Hofman, Mads V. Hollegaard, David M. Hougaard, Masashi Ikeda, Inge Joa, Antonio Julia, Rene S. Kahn, Luba Kalaydjieva, Sena Karachanak-Yankova, Juha Karjalainen, David Kavanagh, Matthew C. Keller, Brian J. Kelly, James L. Kennedy, Andrey Khrunin, Yunjung Kim, Janis Klovins, James A. Knowles, Bettina Konte, Vaidutis Kucinskas, Zita Ausrele Kucinskiene, Hana Kuzelova-Ptackova, Anna K. Kahler, Claudine Laurent, Jimmy Lee Chee Keong, S. Hong Lee, Sophie E. Legge, Bernard Lerer, Miaoxin Li, Tao Li, Kung-Yee Liang, Jeffrey Lieberman, Svetlana Limborska, Carmel M. Loughland, Jan Lubinski, Jouko Lnnqvist, Milan Macek, Jr., Patrik K.E. Magnusson, Brion S. Maher, Wolfgang Maier, Jacques Mallet, Sara Marsal, Manuel Mattheisen, Morten Mattingsdal, Robert W. McCarley, Colm McDonald, Andrew M. McIntosh, Sandra Meier, Carin J. Meijer, Bela Melegh, Ingrid Melle, Raquelle I. Mesholam-Gately, Andres Metspalu, Patricia T. Michie, Lili Milani, Vihra Milanova, Younes Mokrab, Derek W. Morris, Ole Mors, Preben B. Mortensen, Kieran C. Murphy, Robin M. Murray, Inez Myin-Germeys, Bertram Mller-Myhsok, Mari Nelis, Igor Nenadic, Deborah A. Nertney, Gerald Nestadt, Kristin K. Nicodemus, Liene Nikitina-Zake, Laura Nisenbaum, Annelie Nordin, Eadbhard O’Callaghan, Colm O’Dushlaine, F. Anthony O’Neill, Sang-Yun Oh, Ann Olincy, Line Olsen, Jim Van Os, Psychosis Endophenotypes International Consortium, Christos Pantelis, George N. Papadimitriou, Sergi Papiol, Elena Parkhomenko, Michele T. Pato, Tiina Paunio, Milica Pejovic-Milovancevic, Diana O. Perkins, Olli Pietilinen, Jonathan Pimm, Andrew J. Pocklington, John Powell, Alkes Price, Ann E. Pulver, Shaun M. Purcell, Digby Quested, Henrik B. Rasmussen, Abraham Reichenberg, Mark A. Reimers, Alexander L. Richards, Joshua L. Roffman, Panos Roussos, Douglas M. Ruderfer, Veikko Salomaa, Alan R. Sanders, Ulrich Schall, Christian R. Schubert, Thomas G. Schulze, Sibylle G. Schwab, Edward M. Scolnick, Rodney J. Scott, Larry J. Seidman, Jianxin Shi, Engilbert Sigurdsson, Teimuraz Silagadze, Jeremy M. Silverman, Kang Sim, Petr Slominsky, Jordan W. Smoller, Hon-Cheong So, Chris C.A. Spencer, Eli A. Stahl, Hreinn Stefansson, Stacy Steinberg, Elisabeth Stogmann, Richard E. Straub, Eric Strengman, Jana Strohmaier, T. Scott Stroup, Mythily Subramaniam, Jaana Suvisaari, Dragan M. Svrakic, Jin P. Szatkiewicz, Erik Sderman, Srinivas Thirumalai, Draga Toncheva, Paul A. Tooney, Sarah Tosato, Juha Veijola, John Waddington, Dermot Walsh, Dai Wang, Qiang Wang, Bradley T. Webb, Mark Weiser, Dieter B. Wildenauer, Nigel M. Williams, Stephanie Williams, Stephanie H. Witt, Aaron R. Wolen, Emily H.M. Wong, Brandon K. Wormley, Jing Qin Wu, Hualin Simon Xi, Clement C. Zai, Xuebin Zheng, Fritz Zimprich, Naomi R. Wray, Kari Stefansson, Peter M. Visscher, Wellcome Trust Case Control Consortium, Rolf Adolfsson, Ole A. Andreassen, Douglas H.R. Blackwood, Elvira Bramon, Joseph D. Buxbaum, Anders D. Børglum, Sven Cichon, Ariel Darvasi, Enrico Domenici, Hannelore Ehrenreich, Tonu Esko, Pablo V. Gejman, Michael Gill, Hugh Gurling, Christina M. Hultman, Nakao Iwata, Assen V. Jablensky, Erik G. Jonsson, Kenneth S. Kendler, George Kirov, Jo Knight, Todd Lencz, Douglas F. Levinson, Qingqin S. Li, Jianjun Liu, Anil K. Malhotra, Steven A. McCarroll, Andrew McQuillin, Jennifer L. Moran, Preben B. Mortensen, Bryan J. Mowry, Markus M. Nthen, Roel A. Ophoff, Michael J. Owen, Aarno Palotie, Carlos N. Pato, Tracey L. Petryshen, Danielle Posthuma, Marcella Rietschel, Brien P. Riley, Dan Rujescu, Pak C. Sham, Pamela Sklar, David St. Clair, Daniel R. Weinberger, Jens R. Wendland, Thomas Werge, Mark J. Daly, Patrick F. Sullivan, and Michael C. O’Donovan.

## 5 Members of the Discovery, Biology, and Risk of Inherited Variants in Breast Cancer (DRIVE) study

The members of DRIVE are Peter Kraft, David J Hunter, Muriel Adank, Habibul Ahsan, Kristiina Aittomki, Lars Beckman, Sonja Berndt, Carl Blomquist, Federico Canzian, Jenny Chang-Claude, Stephen J. Chanock, Laura Crisponi, Kamila Czene, Norbert Dahmen, Isabel dos Santos Silva, Douglas Easton, A. Heather Eliassen, Jonine Figueroa, Olivia Fletcher, Montserrat Garcia-Closas, Mia M. Gaudet, Lorna Gibson, Christopher A. Haiman, Per Hall, Aditi Hazra, Hereditary Breast and Ovarian Cancer Research Group Netherlands (HEBON), Rebecca Hein, Brian E. Henderson, Albert Hofman, John L. Hopper, Astrid Irwanto, Rudolf Kaaks, Muhammad G. Kibriya, Peter Lichtner, Sara Lindstrm, Jianjun Liu, Enes Makalic, Alfons Meindl, Hanne Meijers-Heijboer, Bertram Mller-Myhsok, Taru A. Muranen, Heli Nevanlinna, Julian Peto, Ross L. Prentice, Nazneen Rahman, Daniel F. Schmidt, Rita K. Schmutzler, Melissa C. Southey, Rulla Tamimi, Clare Turnbull, Andre G. Uitterlinden, Rob B. van der Luijt, Quinten Waisfisz, Zhaoming Wang, Alice S. Whittemore, Rose Yang, Wei Zheng.

